# Orchestrating the Acquisition of Oligodendrocyte Precursor Cell versus Olfactory Bulb Interneuron Fates through *Olig1/2* during Mammalian Cortical Gliogenesis and Gliomagenesis

**DOI:** 10.1101/2024.11.11.623106

**Authors:** Yu Tian, Ziwu Wang, Feihong Yang, Wen Zhang, Jialin Li, Lin Yang, Tongye Fu, Wenhui Zheng, Zhejun Xu, Tong Ma, Yan You, Xiaosu Li, Jiangang Song, Yunli Xie, Zhengang Yang, Zhuangzhi Zhang

## Abstract

The hijacking of developmental gliogenesis programs is a hallmark of glioblastoma (GBM), in which glial precursor cells (GPCs) typically differentiate into both neurons and glial cells. In GBM, this process is disrupted, leading to the overproduction of proliferative glial-like cells. Our study demonstrates that the knockout of *Olig1/2* in both normal development and gliomagenesis causes GPCs to shift from generating highly proliferative oligodendrocyte precursor cells to producing non-proliferative olfactory bulb interneurons. Mechanistically, *Olig1/2* play dual roles by orchestrating distinct transcriptional programs in GPCs, particularly inhibiting the expression of *Gsx2* through direct binding to its multiple enhancers. Additionally, we provide compelling evidence that human H3.3G34R/V-mutant tumors, a subtype of high-grade gliomas, originate from dorsal cortical-derived GPCs rather than from the previously assumed progenitors in the ventral basal ganglia. Collectively, our findings reveal a previously unrecognized role of *Olig1/2* in both gliogenesis and gliomagenesis, offering deeper insights into the connections between normal neural development and tumorigenesis.

**Graphical Abstract:** 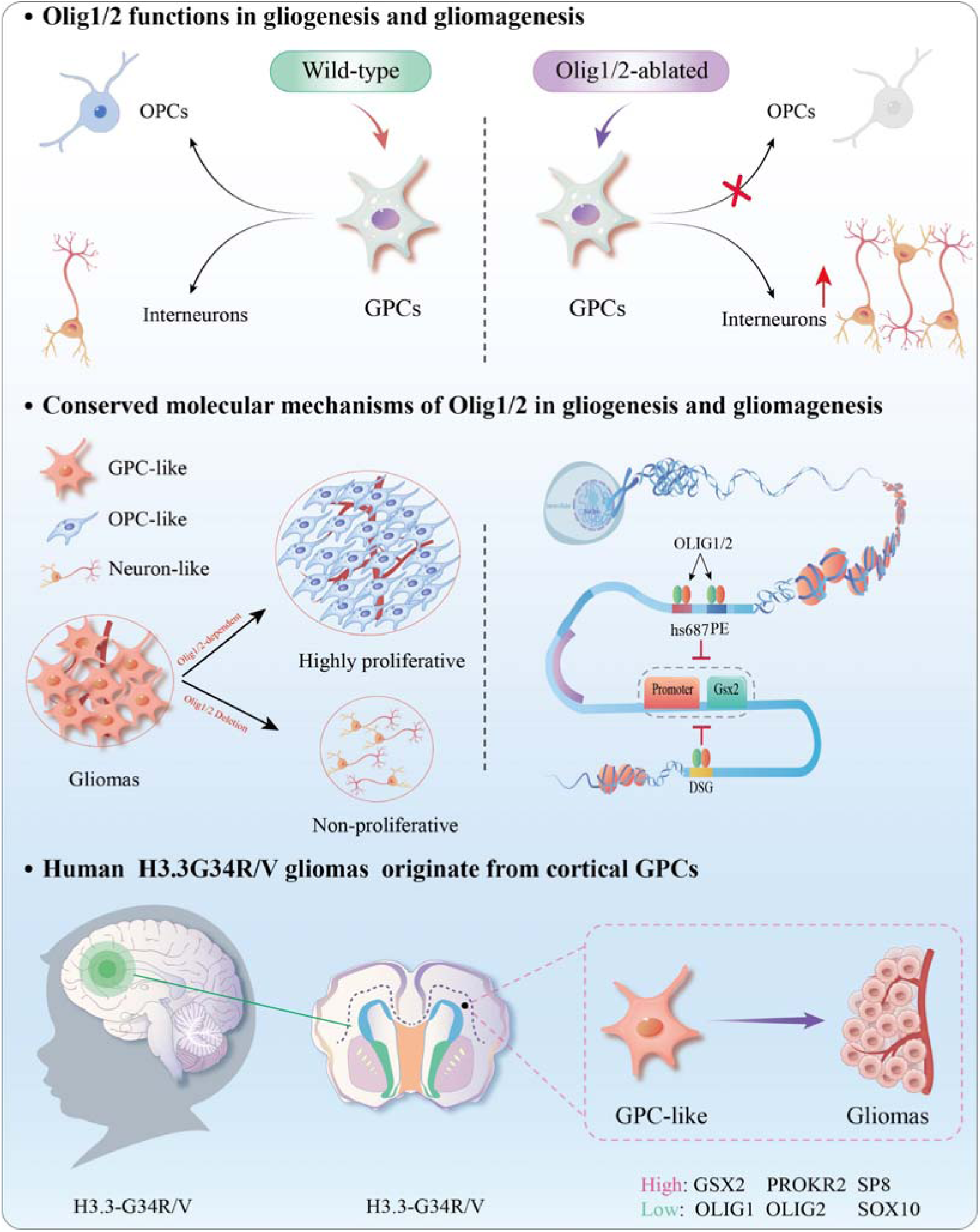

## INTRODUCTION

The human neocortex, vital for human higher-order brain functions like language and cognition, predominantly consists of neurons and glia cells. Glia cells, including astrocytes and oligodendrocytes, play variety of essential roles that support and regulate the function of neurons, ensuring proper brain function^1–3^. These non-neuronal cells are derived from the same glial progenitor cells (GPCs) during gliogenesis^4–8^. In healthy brain, GPCs can also differentiate into neurons under certain conditions, highlighting the plasticity of neural cell fate^9–12^. Key signaling pathways, including *Wnt*, *Notch*, and Sonic Hedgehog (*Shh*), along with transcription factors like *Nfia* and *Oct4*, govern this fate decision^6,13–18^. However, in gliomas, particularly malignant ones like glioblastomas, this differentiation process is disrupted. Mutations and dysregulated signaling pathways cause GPCs^19^ or tumor-initiating cells^20^ to generate oligodendrocyte-like and astrocyte-like cells that continue to proliferate instead of producing neurons ^21–24^. Therefore, studying gliogenesis can help identify key molecular and genetic alterations that cause GPCs and their progeny to undergo uncontrolled proliferation, ultimately leading to tumor formation.

In terms of treatment, inducing the transdifferentiation of glioma cells into neurons is emerging as a promising strategy. Since neurons are post-mitotic and cannot divide, converting tumor cells into neurons could slow tumor progression and reduce malignancy. The bHLH transcription factors (TFs) and their cofactors have been demonstrated to be essential for gliogenesis. *Olig1* and *Olig2*, two bHLH TFs known for their critical roles in oligodendrocyte (OL) development^25–33^, are highly expressed in GPCs. Additionally, these two genes are markedly overexpressed in proneural-like glioblastoma (GBM), one of the most lethal forms of glioma, with a median survival of less than one year after diagnosis^24,34,35^. Although previous studies have shown that *Olig1* and *Olig2* share high homology in their structural domains and exhibit coordinated expression in the CNS^36^, their roles in GPCs, as well as the relationship between these two genes in cortical gliogenesis and gliomagenesis remian poorly understood.

Different from adult GBM, histone H3.3G34R/V gliomas, a subtype of pediatric high-grade gliomas, exhibit both neuronal and astroglial identities while lacking oligodendroglial identity ^21,37–39^. These gliomas typically occur in the cerebral hemispheres, one of its characteristics is the low expression levels of *OLIG1* and *OLIG2* and their transcriptomic profile suggests an association with early forebrain development^21,38–41^. A groundbreaking study proposes that tumor-initiating cells of histone H3.3G34R/V glioma may derive from GSX2-expressing cells located in the lateral ganglionic eminence (LGE)^42,43^. This conclusion is based on the exclusive expression of the *GSX2* in ganglionic eminence (GE)-derived progenitors. Unlike previous studies, our recent findings reveal that cortical-derived progenitors also express *Gsx2* ^6,7,44^. Therefore, we aim to determine whether H3.3G34R/V glioma also originate in cortical GSX2-expressing GPCs.

In this study, we generated multiple *Olig* mutant mouse models to investigate the role of *Olig1/2* in glial progenitor cells (GPCs) during gliogenesis and glioma development. Our findings demonstrate, for the first time, that the loss of *Olig1/2* results in GPCs failing to produce highly proliferative oligodendrocyte precursor cells (OPCs), instead driving their differentiation into OB interneurons (INs) with limited proliferative potential. By integrating multiple omics technologies, including scRNA-Seq, CUT&Tag-Seq, and scATAC-Seq, we discovered that *Olig1*/*2* cooperatively promote OPC generation while suppressing OB interneuron production by repressing *Gsx2* expression in GPCs. More importantly, we identified that this molecular mechanism is also involved in gliomagenesis. Additionally, we found that cortical GPCs, rather than ventral ganglionic eminence progenitors, are the key tumor-initiating cells in human H3.3G34R/V gliomas. Therefore, those findings will deepen our understanding the gaps between normal development and tumorigenesis.

## RESULTS

### 1. Increased cortical-derived olfactory bulb (OB) interneuron production in *Olig1/2* double-mutant mice

During gliogenesis and gliomagenesis, bHLH transcription factors *Olig1/2* are highly expressed in glial progenitor cells (GPCs), which have the potential to generate both glial cells and neurons^2,6,7,19^. However, the specific functions of these two transcription factors in GPCs remain unclear. To systematically investigate the roles of *Olig1*/*2* in cortical GPCs, we generated various *Olig* mutant mice, including *Olig1* conventional knockout (*Olig1*-KO) mice, *Olig2* conditional knockout (*Olig2*-CKO) mice, and *Olig1/2* double conditional knockout (*Olig1/2*-CKO) mice (Figure S1a). To explore the function of *Olig1/2* in the cortex, we cross a *Emx1-Cre* mouse line with *Olig2^F/F^* or *Olig1/2^F/F^* mouse line to obtain the *Emx1-Cre; Olig2^F/F^* (*Olig2*-ECKO) or *Emx1-Cre; Olig1/2^F/F^* (*Olig1/2*-ECKO) mice. First, we analyzed the expression of the OB interneuron marker genes, such as DLX2 and SP8 via immunostaining. Our results showed that a large number of the OB interneurons were accumulated in the dorsal subventricular zone (dSVZ) of the *Olig1/2*-ECKO mice, while there were no significant changes in the *Olig1*-KO and *Olig2*-ECKO mice at P0, compared to control mice (Figure 1b and Figure S1b). However, we observed a significant increase in the number of OB interneurons in the dSVZ of *Olig1*-KO, *Olig2*-ECKO and *Olig1/2*-ECKO mice at P7, compared to control mice (Figure 1a). Next, we also analyzed the sagittal sections, which revealed that DLX2-positive cells were significantly increased in the dSVZ of *Olig1*-KO, *Olig2*-ECKO and *Olig1/2*-ECKO mice at P7, compared to control mice (Figure S1c). The number of the SP8-positive cells are also increased in the OB of *Olig1/2*-ECKO mice, compared to control mice (Figure S1d). Even in adult stage, we observed a significant increase in the number of SP8- or DLX2-positive neurons in the *Olig1/2*-ECKO mice (Figure 1b and Figure S1e). Notably, the number of the OB interneurons significantly increased in the *Olig1/2*-ECKO mice from P0 to P30 compared to the single mutant and wild-type mice (Figure 1b).

**Fig. 1.**
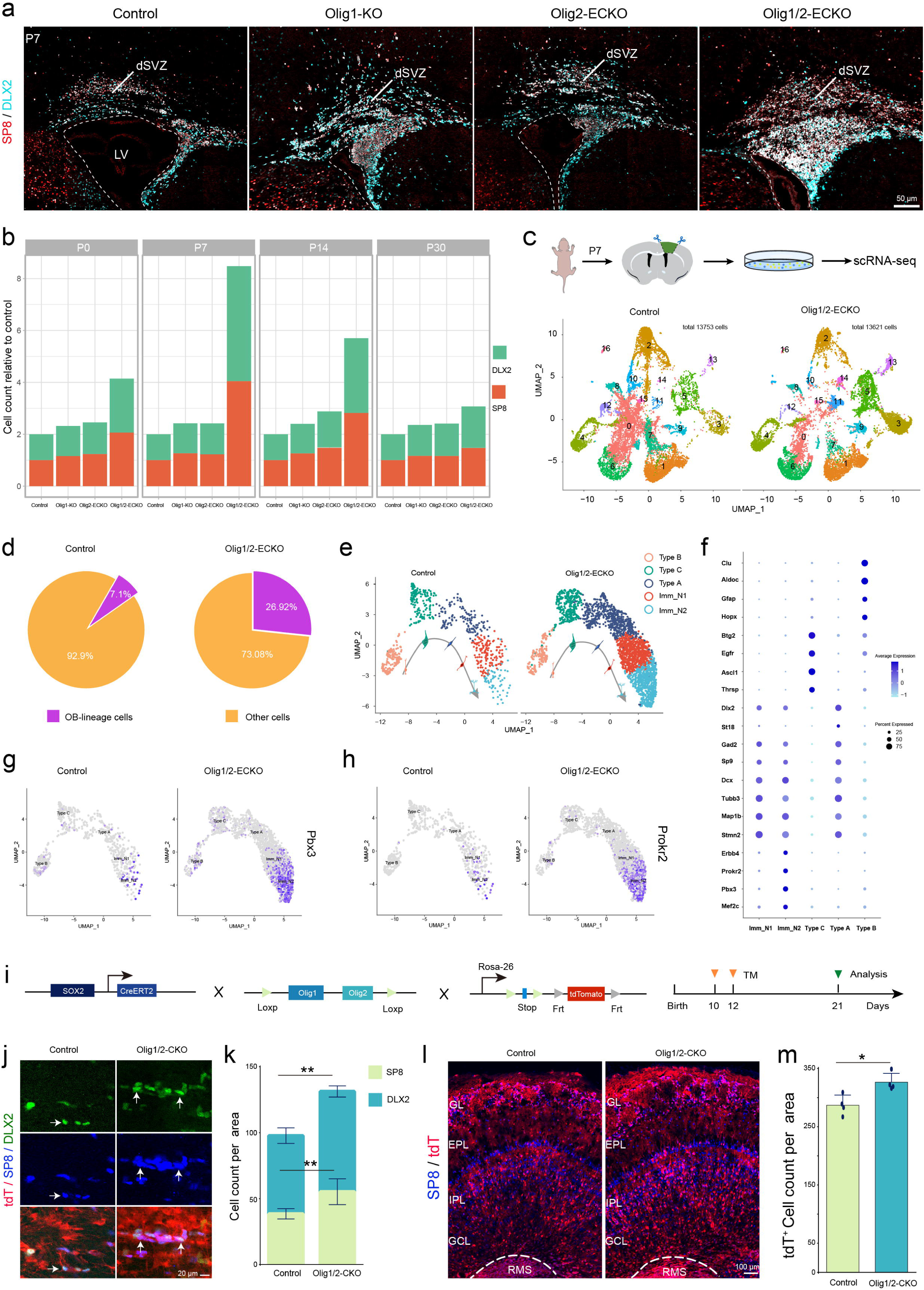
*Olig1* and *Olig2* work together to inhibit the generation of OB interneurons. (a) The number of the SP8- and DLX2-positive cells are significantly increased in dSVZ of the *Olig1*-KO, *Olig2*-ECKO and *Olig1/2*-ECKO mice, compared to control mice at P7. Notably, *Olig1/2*-ECKO mice exhibit a higher number of SP8- and DLX2-positive cells than the other genotypes. (b) The statistics show that the SP8- and DLX2-positive cells are increased in *Olig1/2*-ECKO mice from P0 to P30, compared to control mice. The number of the OB interneurons in *Olig1*-KO and *Olig2*-ECKO mice are slightly increased from P7 to P30, compared to control mice. (c) The scRNA-seq data of the wild-type and *Olig1/2*-ECKO mice cortex at P7. (d) The proportion of OB-lineage cells is increased in the *Olig1/2*-ECKO mice, compared to control mice. (e) The OB-lineage cells are further divided in to five clusters, including Type B, Type C, Type A, Immature interneuron 1(Imm_N1) and Immature interneuron 2 (Imm_N2). (f) The bubble chart showing the differentially expressed genes (DEGs) between five clusters. (g-h) The expression of the Pbx3 and Prokr2 in *Olig1/2*-ECKO and control mice. (i) The process diagram for experimental design. (j) The SP8/DLX2/tdT triple immunostaining between Sox2-CreER; *Olig1/2^F/F^*; *IS^F/+^* (*Olig1/2*-CKO) and control mice. (k) The SP8/tdT and DLX2/tdT double-positive cells are increased in *Olig1/2*-CKO mice, compared to control mice. (l) The immunostaining showing SP8/tdT double-positive cells in the OB. (m) Statistical data showing SP8/tdT double-positive cells are increased in OB of the *Olig1/2*-CKO mice, compared to control mice. *P<0.05, **P<0.01 and ***P<0.001 (one-way ANOVA followed by Tukey–Kramer post-hoc test). N=4 mice per group, mean±s.e.m. SVZ, subventricular zone; GL, glomerular layer; EPL, external plexiform layer; GCL, granule cell layer; IPL, internal plexiform layer. LV, Lateral ventricle, RMS, rostral migratory stream.

Second, we collected the cortex of control and *Olig1/2*-ECKO mice for scRNA-seq at P7. After removing low-quality cells, we obtain 13753 cells from control mice and 13621 cells from *Olig1/2*-ECKO mice. UMAP was performed on these cells using Seurat 3.2, resulting in 17 clusters (C0–C16). Using marker genes to define these clusters more precisely, we identified 10 discrete populations: endothelial cells (C0 and C6), astrocytes (C1 and C7), OL-lineage cells (C2 and C10), OB-lineage cells (C3 and C5), immune cells (C4 and C12), ependymal cells (C8), mural cells (C9), PyN-lineage cells (C11, C13 and C15), erythrocyte cells (C14), leukocyte cells (C16) (Figure 1c and Table 1). In the control mice, OB-lineage cells accounted for approximately 7.1%, whereas in the *Olig1/2*-ECKO mice, this proportion was approximately 26.92% (Figure 1d), suggesting that *Olig1/2* ablation leads to increased production of OB interneurons. To further analyze whether OB-lineage cells can develop normally after the loss of *Olig1/2*, we extracted OB-lineage cells and divided them into five groups: Type B (expressing *Clu*, *Gfap*, and *Hopx*), Type C (expressing *Btg2*, *Ascl1*, and *Egfr*), Type A (expressing *Dlx2*, *Gad1*, and *St18*), immature N1 (expressing *Dcx*, *Tubb3*, and *Map1b*), and immature N2 (expressing *Erbb4*, *Prokr2*, and *Pbx3*) (Figures 1e-f). We found that these *Olig1/2*-ablated OB interneurons could express *Prokr2* and *Pbx3*, which are molecular markers representing OB interneuron migration (Figures 1g-h). Therefore, the loss of *Olig1/2* leads to an increase in OB interneurons without affecting their normal migration process.

Finally, the previous studies showed that *Olig1/2* are also expressed in Type C cells during adult neurogenesis^45^. To investigate the roles of *Olig1/2* in this process, we crossed a *Sox2-CreER* mouse line with *Olig1/2^F/F^; IS^F/+^* reporter mice, ultimately obtaining *Sox2-CreER; Olig1/2^F/F^; IS^F/+^* mice. We administered tamoxifen (TM) injections to the mice at P10 and P12, and conducted analysis at P21 (Figure 1i). Our results showed that tdT/SP8 or tdT/DLX2 double-positive cells are increased in the *Sox2-CreER; Olig1/2^F/F^; IS^F/+^* (*Olig1/2*-CKO) mice, compared to control mice (Figures 1j-k). Even in OB, the tdT/SP8 double-positive cells are also increased (Figures 1l-m). Taken together, our findings indicate that *Olig1/2* coordinately inhibit the production of OB interneuron from embryonic to postnatal stages.

### 2. Functional similarities and differences between *Olig1* and *Olig2* in driving OPC production and myelination

Previous studies have shown that *Olig1/2* are crucial for the development of oligodendrocytes, but there is controversy regarding the roles of *Olig1* and *Olig2* in the generation of oligodendrocyte progenitor cells and the process of cortical myelination^27–29,32,33,46–48^. To investigate the function of the *Olig1/2* in cortical-derived OL-lineage cells, we first explore the expression of *Sox10* in *Olig* mutant mice. Our results showed that the number of the SOX10-positive cells significantly decreased in the *Olig2*-ECKO and *Olig1/2*-ECKO mice at P0, while only slightly decreased in *Olig1*-KO mice compared to control mice (Figures 2a-b). Indeed, the number of SOX10-positive cells continuously decreased from P0 to P30 in *Olig1/2*-ECKO mice (Figure 2c). These results indicate that *Olig1* plays a limited role in the generation of cortical OPCs, while *Olig2* plays a critical role in this process. It is worth noting that these SOX10-positive cells expressed OLIG2, suggesting that those cells are not cortical-derived. Our recent studies found that cortical OL-lineage cells are derived from MGE and cortical RGCs and OL-lineage cells derived from the MGE are generated and mature earlier^44^. During the development of OLs, they sequentially express *Sox10*, *S100b*, and *Mbp*^49^ (Figure S2c). We found that more than 90 percent of SOX10/OLIG2 double-positive cells in the *Olig2*-ECKO and *Olig1/2*-ECKO mice are also expressed S100B, suggesting that these cells are MGE-derived (Figures S2a-b). Additionally, we analyzed scRNA-Seq from P7 mice mentioned earlier. We found that *Sox10* is primarily expressed in cluster 2, which corresponds to OL-lineage cells (Figure S2d). The differential expression gene analysis between the *Olig1/2*-ECKO and control groups showed that OL-lineage cells in *Olig1/2*-ECKO mice exhibited higher expression of *Nkx2-1* and *S100b*, and lower expression of *Emx1*, compared to control mice (Figures 2d-e and Figures S2e-f). These findings suggest that the SOX10-positive cells in the *Olig1/2*-ECKO mice are derived from the MGE. Altogether, our results confirm that GPCs rely more on *Olig2* than *Olig1* to generate OPCs.

**Fig. 2.**
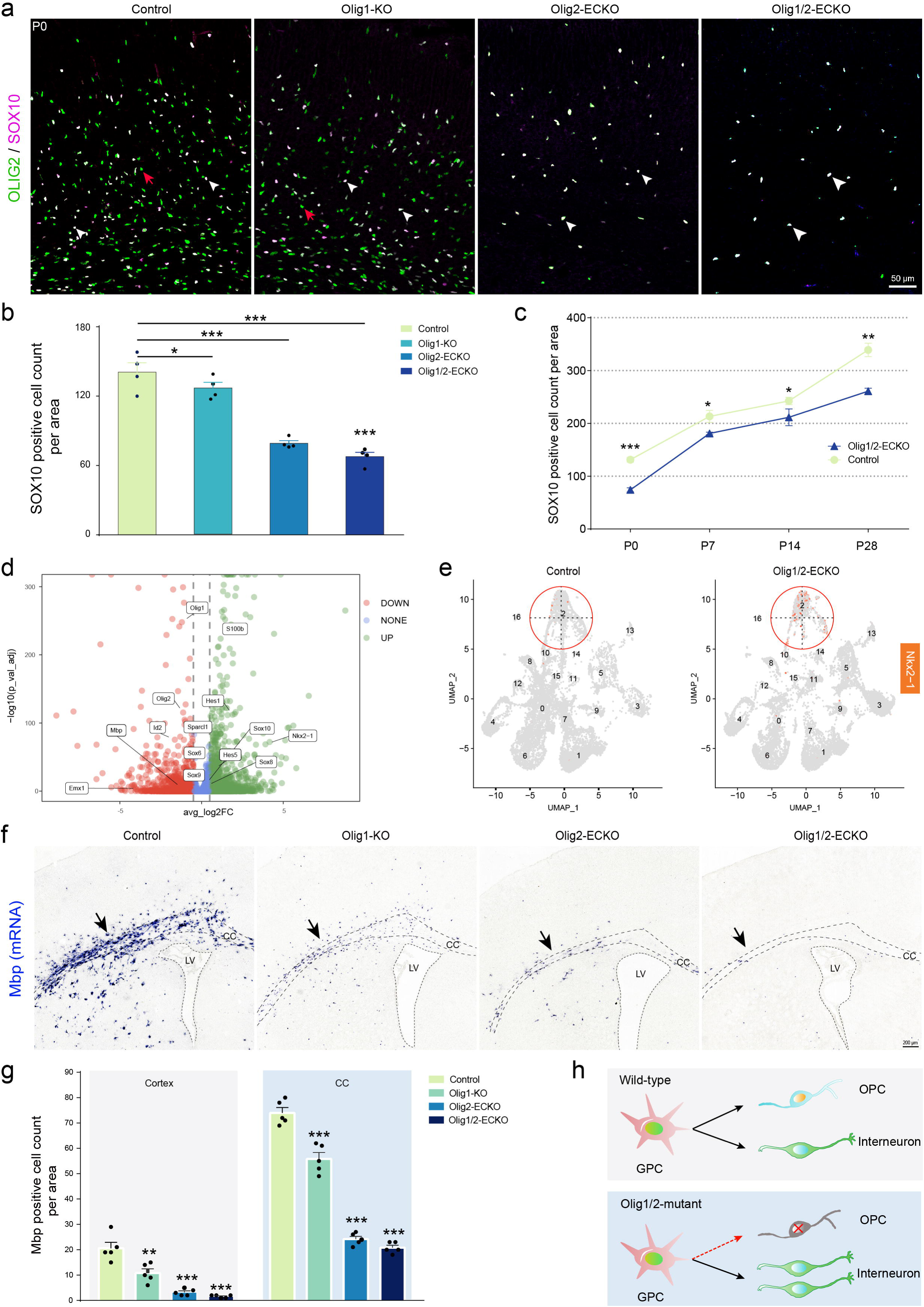
*Olig1/2* are required for cortical OPC generation and myelination. (a) The immunostaining of the OLIG2 and SOX10 in the cortex of control, *Olig1*-KO, *Olig2*-ECKO and *Olig1/2*-ECKO mice at P0. (b) The histogram shows the SOX10-positive cell count in the cortex of the control, *Olig1*-KO, *Olig2*-ECKO and *Olig1/2*-ECKO mice at P0. (c) The SOX10-positive cell count in the *Olig1/2*-ECKO mice is consistent reduced from P0 to P30, compared to control mice. (d) The volcano plot shows the DEGs of OL-lineage cells in the *Olig1/2*-ECKO mice at P7, compared to control mice. (e) The expression level of the *Nkx2-1* in *Olig1/2*-ECKO mice is higher than that in control mice at P7. (f) *In situ* hybridization shows that *Mbp* expression is significantly reduced in the *Olig1*-KO, *Olig2*-ECKO and *Olig1*/*2*-ECKO mice at P7, compared to control mice. (g) The statistics show that the *Mbp* expression is significantly reduced in the *Olig1*-KO, *Olig2*-ECKO and *Olig1*/*2*-ECKO mice at P7, both in cortex and corpus callosum (CC), compared to control mice. **P<0.01 and ***P<0.001 (one-way ANOVA followed by Tukey–Kramer post-hoc test). N=4 mice per group, mean±s.e.m. (h) A potential model of the *Olig1*/*2* function in the tri-IPC. LV, Lateral ventricle.

To investigate the roles of *Olig1/2* in the myelination process, we performed in situ hybridization to assess changes in *Mbp* expression^50,51^, a gene critical for the formation and maintenance of the myelin sheath. Our results show that the expression of the *Mbp* is significantly decreased in both the cortex and corpus callosum of *Olig1*-KO, *Olig2*-ECKO and *Olig1/2*-ECKO mice at P7, compared to control mice (Figures 2f-g). In *Olig2*-ECKO and *Olig1/2*-ECKO mice, the reduction of *Mbp* expression was more pronounced (Figure 2f). In adulthood, myelination partially recovered in *Olig1*-KO, *Olig2*-ECKO and *Olig1/2*-ECKO mice compared to control mice, but it did not fully return to normal levels (Figures S2g-h). Therefore, our results suggest that *Olig1* and *Olig2* have similar functions in cortical myelination. Combining the above results, the loss of *Olig1/2* in GPCs leads to an increase in OB neurons and the elimination of cortical-derived OPCs during gliogenesis (Figure 2h).

### 3. scRNA-Seq reveals that GPCs transit from producing OPCs to OB interneurons in *Olig1/2* double-mutant mice

To investigate whether the increase of OB interneurons following *Olig1/2* ablation is due to fate change in GPCs, we performed in utero electroporation (IUE) to deliver *Cre* plasmids specifically to cortical RGCs of *IS^F/+^* or *Olig1/2^F/F^; IS^F/+^* mice. FACS was then used to sort *Cre*-recombined cells for scRNA-seq (Figure 3a). After obtaining high-quality cells, we performed clustering analysis and identified seven cell clusters (Figure 3b and Table 2). The glutaminergic projection pyramidal neurons (PyNs) express high level of *Neurod1*, *Neurod6* and *Bhlhe22*; the olfactory bulb (OB) interneurons express high level of *Dlx1*, *Nrxn3* and *Tshz1*; the RGCs express high level of *Slco1c1*, *Sparc* and *Gfap*; the astrocytes (As) express high level of *Aldh1l1*, *Tnc* and *Id1*; the GPCs express high level of *Ascl1*, *Egfr* and *Rrm2*; the ependymal cells express high level of *Foxj1*, *S100a11* and *Ogn*; and the OPCs express high level of *Gpr17*, *Cspg4* and *Martn4* (Figure 3c and Table 2). We found that the OPCs in *Olig1/2*-CKO mice were significantly reduced compared to the control mice (Figure 3d). Although a small number of cells in the *Olig1/2*-CKO mice expressed OPC genes such as *Pdgfra*, *Sox10*, and *Cntn1*, we found that these cells also expressed *Olig1* and *Olig2* (Figure 3d and Figures S3a-b). These results suggest that these OPCs might be incompletely *Cre*-recombined cells or a small fraction of tdT-negative cells mixed during the FACS sorting process. On the other hand, we observed a significant increase in OB interneurons in *Olig1/2*-CKO mice; those cells expressed *Dlx2*, *Sp9* and *Gad2* (Figure 3d and Figure S3c).

**Fig. 3.**
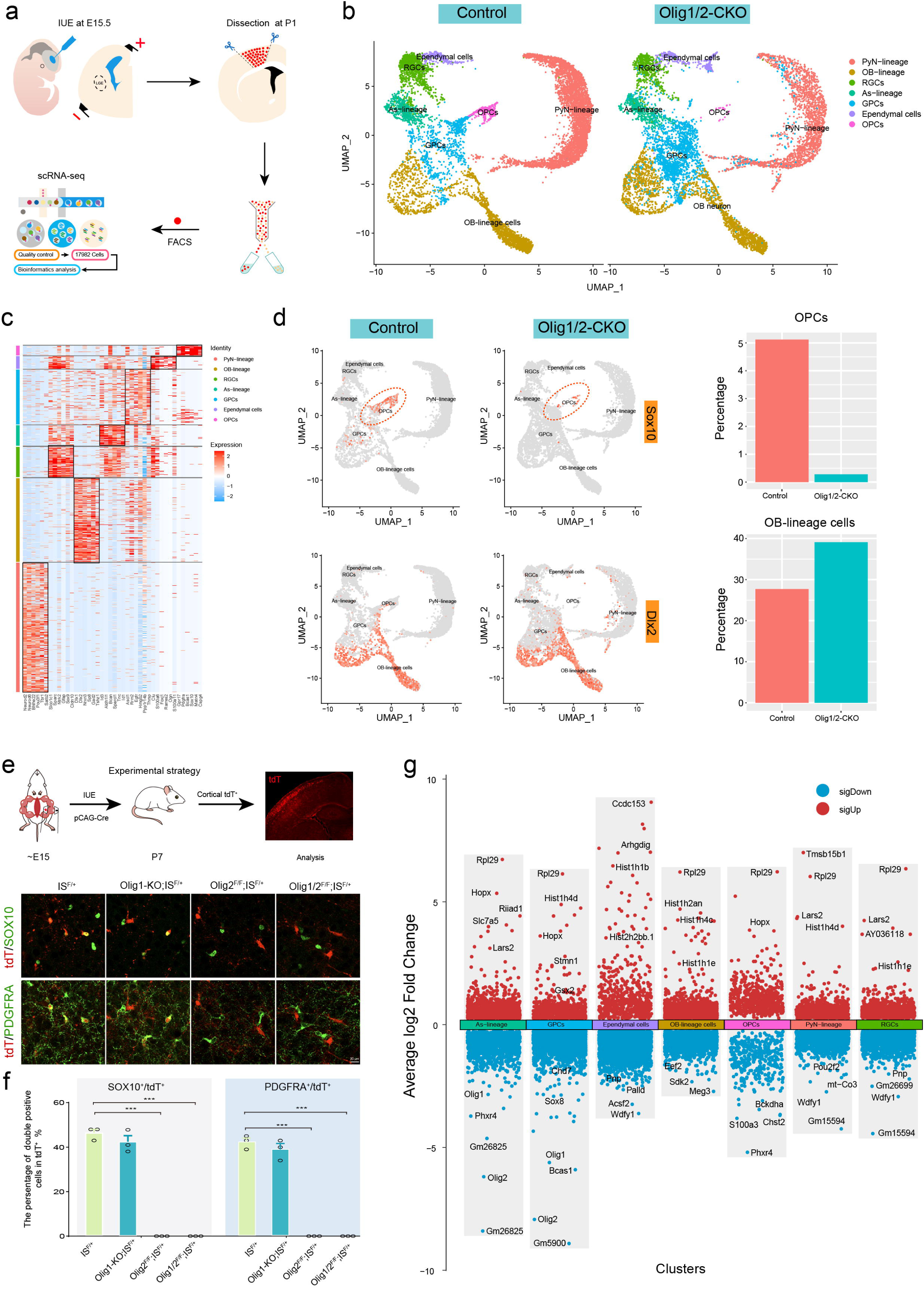
Fate switching of GPCs from OPCs into OB INs in the *Olig1/2*-CKO mice. (a) The diagram of scRNA-Seq experiment workflow. (b) Dividing the P1 scRNA-Seq into 7 clusters, including PyN-lineage cells, OB-lineage cells, RGCs, As-lineage cells, GPCs, Ependymal cells and OPCs. (c) Heatmap shows the differentially expressed genes in 7 clusters. (d) The OL-lineage cells, which expressed *Sox10* are significantly decreased, whereas the OB-lineage cells, which expressed *Dlx2* are significantly increased in the *Olig1/2*-CKO mice compared to control mice. (e) Combining in utero electroporation (IUE) with immunohistochemistry to show tdT/SOX10 double-positive cells or tdT/PDGFRA double-positive cells in the cortex of the *IS^F/+^*, *Olig1*-KO*; IS^F/+^*, *Olig2^F/F^; IS^F/+^* and *Olig1/2^F/F^; IS^F/+^* mice at P7. (f) The statistical data shows that tdT/SOX10 double-positive cells or tdT/PDGFRA double-positive cells are no longer generated in the *Olig2^F/F^; IS^F/+^* and *Olig1/2^F/F^; IS^F/+^* mice. ***P<0.001 (one-way ANOVA followed by Tukey–Kramer post-hoc test). N=3 mice per group, mean±s.e.m. (g) Multiple-group volcano plot showing differentially expressed genes of 7 cell clusters between control and *Olig1/2*-CKO mice.

Using immunohistochemistry to validate the scRNA-Seq results in vivo, we found that the number of tdT/PDGFRA and tdT/SOX10 double-positive cells were significantly decreased in the *Olig2^F/F^; IS^F/+^* and *Olig1/2^F/F^; IS^F/+^* mice at P7, compared to control and *Olig1*-KO; *IS^F/+^* mice (Figures 3e-f). In contrast, the number of tdT/SP8 double-positive cells were significantly increased in the *Olig1*-KO; *IS^F/+^*, *Olig2^F/F^; IS^F/+^*and *Olig1/2^F/F^; IS^F/+^* mice, compared to control mice (Figures S3e-f). These findings indicate that *Olig1/2* promote OPC production and simultaneously suppress OB interneuron production in GPCs. Additionally, we also noted an increase in GPCs that highly expressed *Egfr* and *Ascl1* (Figure S3d). The differentially expressed genes analysis of GPCs revealed a significant decrease in the expression of genes associated with OPC fate determination (including *Bcas1*, *Chd7* and *Sox10*) and a significant increase in the expression of genes (including *Stmn1*, *Hist1h4d* and *Gsx2*) associated with OB interneuron fate determination (Figure 3g). Interestingly, we also observed that the generation of As-lineage cells was not affected in *Olig1/2*-CKO mice, compared to control mice (Figure 3b). Therefore, our findings suggest that *Olig1/2* plays a dual role in GPCs, promoting OPC production while inhibiting OB interneuron production in a cell-autonomous manner.

### 4. *Olig1/2* plays dual roles by directing distinct transcriptional programs to regulate GPC fate selection

To investigate the molecular mechanisms by which *Olig1* and *Olig2* regulate OPC and OB interneuron fate determination during gliogenesis, we performed CUT&Tag-Seq and scATAC-Seq^7^ on FACS-sorted GFP^+^ cells from the cortex of *hGFAP*-GFP animals at perinatal stages, when GPCs undergo proliferation and differentiation (Figure 4a). The CUT&Tag-Seq showed high reproducibility, and OLIG1 and OLIG2 binding peaks were enriched at TSS regions, consistent with the characteristics of transcription factors regulating gene expression (Figure 4b and Figures S4a-b). We found that most of the genes bound by OLIG1 are also bound by OLIG2 (Figure 4c). Indeed, motif analysis of binding sites reveals that OLIG1 and OLIG2 shared the similar binding motifs (Figure 4d and Figure S4c). Most of these motifs are typically recognized in bHLH transcription factors, such as *Neurog2*, *Sox10* and *Sox3* (Figure 4d). These results indicate that OLIG1 and OLIG2 co-regulate certain genes to control the cell fate determination during brain development.

**Fig. 4.**
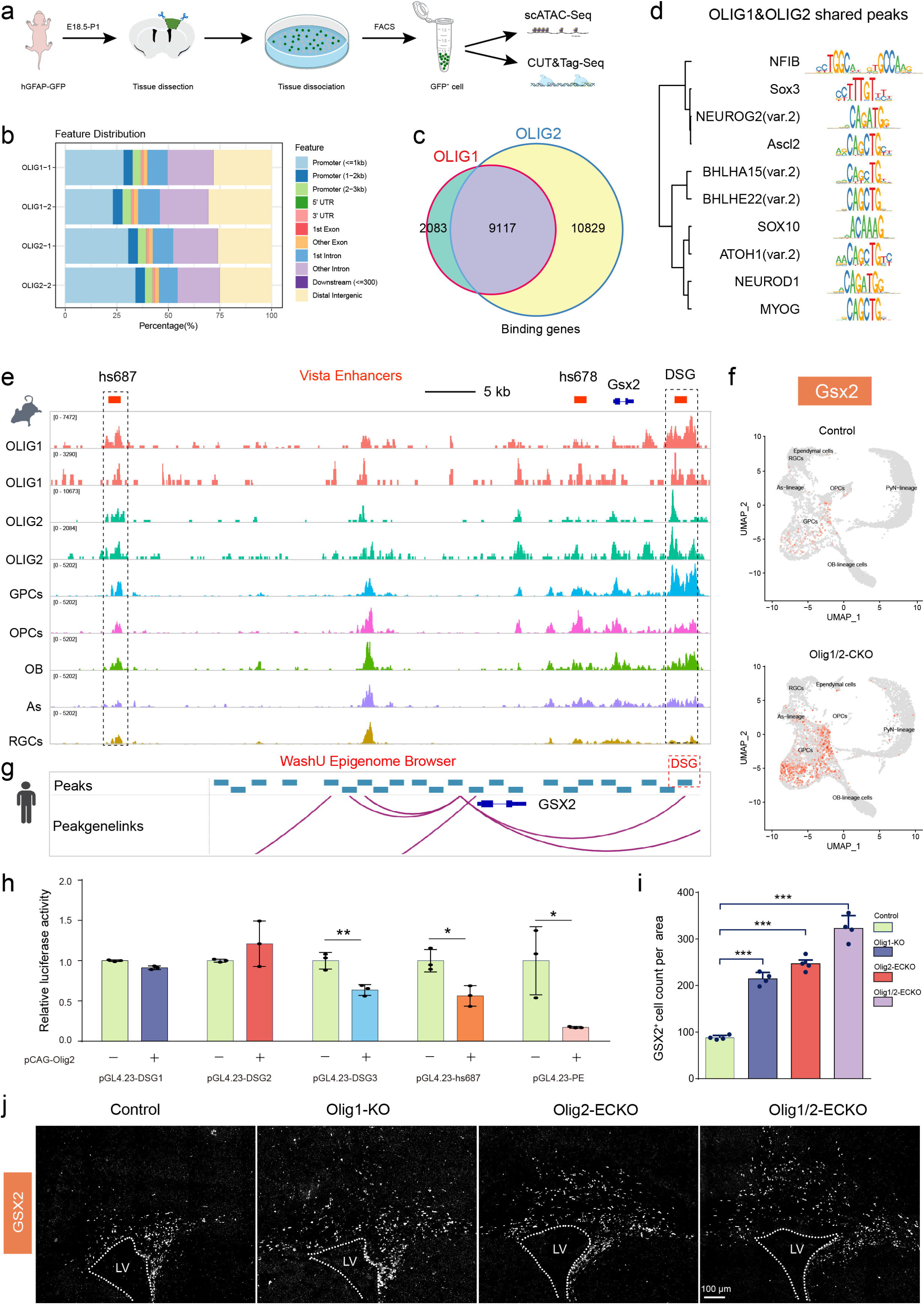
*Olig1/2* inhibit the generation of the OB interneurons by repressing *Gsx2* expression. (a) Using fluorescence-activated cell sorting (FACS) to isolate hGFAP-GFP mouse cortical GFP-positive cells for scATAC-Seq and CUT&Tag-Seq. (b) The distribution of binding sites of the OLIG1 and OLIG2 across the genome. (c) Venn Diagram shows that OLIG1 and OLIG2 co-regulate the expression of a large number of genes. (d) The binding motif that are shared by OLIG1 and OLIG2. (e) Joint analysis of CUT&Tag-Seq and scATAC-Seq suggests that OLIG1 and OLIG2 cooperatively regulate the expression of *Gsx2* in GPCs by binding to multiple enhancers of the *Gsx2* gene. (f) *Gsx2* expression in the scRNA-Seq between control and *Olig1/2*-CKO mice. (g) Hi-C data shows DSG enhancer is also regulating the *GSX2* expression in the human cortex. Wavy purple lines represent predicted interactions between enhancers and the *GSX2* promoter. (h) The dual-luciferase assay showing the activation of OLIG2 at the *Gsx2* enhancers. *P<0.05, and **P<0.01 (one-way ANOVA followed by Tukey–Kramer post-hoc test). N=4 mice per group, mean±s.e.m. (i) GSX2 expression is significantly increased in the dSVZ of the *Olig1*-KO, *Olig2*-ECKO and *Olig1/2*-ECKO mice at P7, compared to control mice. (j) Immunofluorescence staining shows the expression of the GSX2 in control, *Olig1*-KO, *Olig2*-ECKO and *Olig1/2*-ECKO mice at P7. ***P<0.001 (one-way ANOVA followed by Tukey–Kramer post-hoc test). N=3, mean±s.e.m. DSG, downstream from *Gsx2*. PE, potential enhancer.

Previous studies have shown that *Olig2* regulates the expression of the *Pdgfra* and *Sox10* in OL-lineage cells^31,52,53^, but whether *Olig1* regulates these genes remains unclear. Here, we found that both OLIG1 and OLIG2 are directly bind the promoter region of *Pdgfra* (Figure S4d). A putative enhancer of *Pdgfra* is also bound by OLIG1 and OLIG2 (Figure S4d). Notably, the OLIG1 and OLIG2 binding regions show higher chromatin accessibility in GPC and OPC clusters, while in other clusters such as OB-lineage cells, As-lineage cells, and RGCs, these regions are relatively less chromatin accessible based on our scATAC-Seq date. The HMG-domain-containing transcription factors, such as *Sox10* and *Sox8*, are essential for the CNS myelination and myelin maintenance^50,54–56^. Our results showed that OLIG1 and OLIG2 directly bind these genes (Figures S4e-f). According to our scATAC-Seq data, we observed the enhancers U2 and U3 of *Sox10* show higher chromatin accessibility in GPC and OPC clusters than in other clusters (Figure S4f). These results indicate that OLIG1 and OLIG2 together directly regulated a series of genes such as *Pdgfra*, *Sox8* and *Sox10* to control the OL-lineage cell generation and maturation.

Our previous studies have shown that *Gsx1/2-Dlx1/2-Sp8/9* is the key transcription factor regulation pathway controlling the generation and specification of the OB-lineage cells^57–59^. We found that that OLIG1 and OLIG2 directly bind to known enhancers of *Gsx2*, such as hs687 and DSG (downstream from *Gsx2*). Notably, the binding sites of the OLIG1 and OLIG2 show higher chromatin accessibility in the GPC, OPC and OB clusters than in As and RGCs (Figure 4e). We also found that OLIG1 and OLIG2 bind a potential enhancer (PE) near the *Gsx2* gene. This binding site shows higher chromatin accessibility in the GPC cluster than in other clusters (Figure S4h). We then performed a luciferase reporter assay in 293T cells. Co-transfection of cells with an *Olig2* expression vector (pCAG-*Olig2*) and an enhancer-reporter construct (pGL4.23-DSG1, DSG2, DSG3, hs687 or PE) resulted in decreased luciferase activity in pGL4.23-DSG3, pGL4.23-hs687 and pGL4.23-PE compared to the pGL4.23 empty control (Figure 4h). More importantly, according to Hi-C data from Greenleaf’s lab in the WashU Epigenome Browser^60^, we found that the DNA sequence homologous to the mouse DSG sequence in humans is associated with the *GSX2* promoter (Figure 4g). These results indicate that the DSG enhancer promotes *GSX2* expression and it is conserved in the human cortex.

Next, reanalysis of scRNA-Seq data revealed that the expression of the *Gsx2* is significantly increased in the *Olig1/2*-CKO mice compared to control mice at P1 (Figure 4f and Figure S4g). Finally, using immunohistochemistry to further verify the impact of *Olig1* and *Olig2* mutations on *Gsx2* expression in vivo, our results showed that a significant increase in the number of GSX2-positive cells in the dSVZ of the *Olig1*-KO, *Olig2*-ECKO, and *Olig1/2*-ECKO mice compared to control mice at P7 (Figures 4i-j). Altogether, our findings strongly support that *Olig1* and *Olig2* cooperate to inhibit OB interneuron production by regulating *Gsx2* expression in both mice and humans.

### 5. Conserved roles of *Olig1/2* in both gliogenesis and gliomagenesis

The hijacking of early developmental programs is a hallmark of gliomas, where tumor cells resemble neurodevelopmental lineages and exhibit mechanisms of neural stem cell resilience. Glioblastoma (GBM), the most prevalent and malignant brain tumor, is notoriously resistant to conventional therapies^61^. *Olig1/2* is present in various diffuse gliomas, and it is particularly enriched in the proneural (OL signature) GBM subtype. Therefore, we further investigated the roles of *Olig1*/*2* in glioblastoma cells. First, we delivered three plasmids containing PBase, PB-*Pdgfb-dn-P53*, and *Cre* into the cortical RGCs of *IS^F/+^* mice, or *Olig1/2^F/F^; IS^F/+^* mice to construct proneural GBM-like model at P0 by IUE (Figure 5a). We found that the *IS^F/+^* mouse has abnormal brain structure, whereas the *Olig1/2^F/F^; IS^F/+^* mice had normal brain structures, without tumor expansion (Figure 5b). To systematically study the roles of *Olig1* and *Olig2* in glioblastoma, we applied the same method to *IS^F/+^* mice, *Olig1*-KO; *IS^F/+^* mice, *Olig2^F/F^; IS^F/+^* mice, and *Olig1/2^F/F^; IS^F/+^* mice. Our results showed that in the *Olig1*-ablated glioblastoma model, the survival time of the mice was not significantly altered, and they died before P50. Excitingly, the *Olig2*-ablated and *Olig1/2*-ablated glioblastoma mice had significantly extended survival times, with some surviving for more than one year (Figure 5c).

**Fig. 5.**
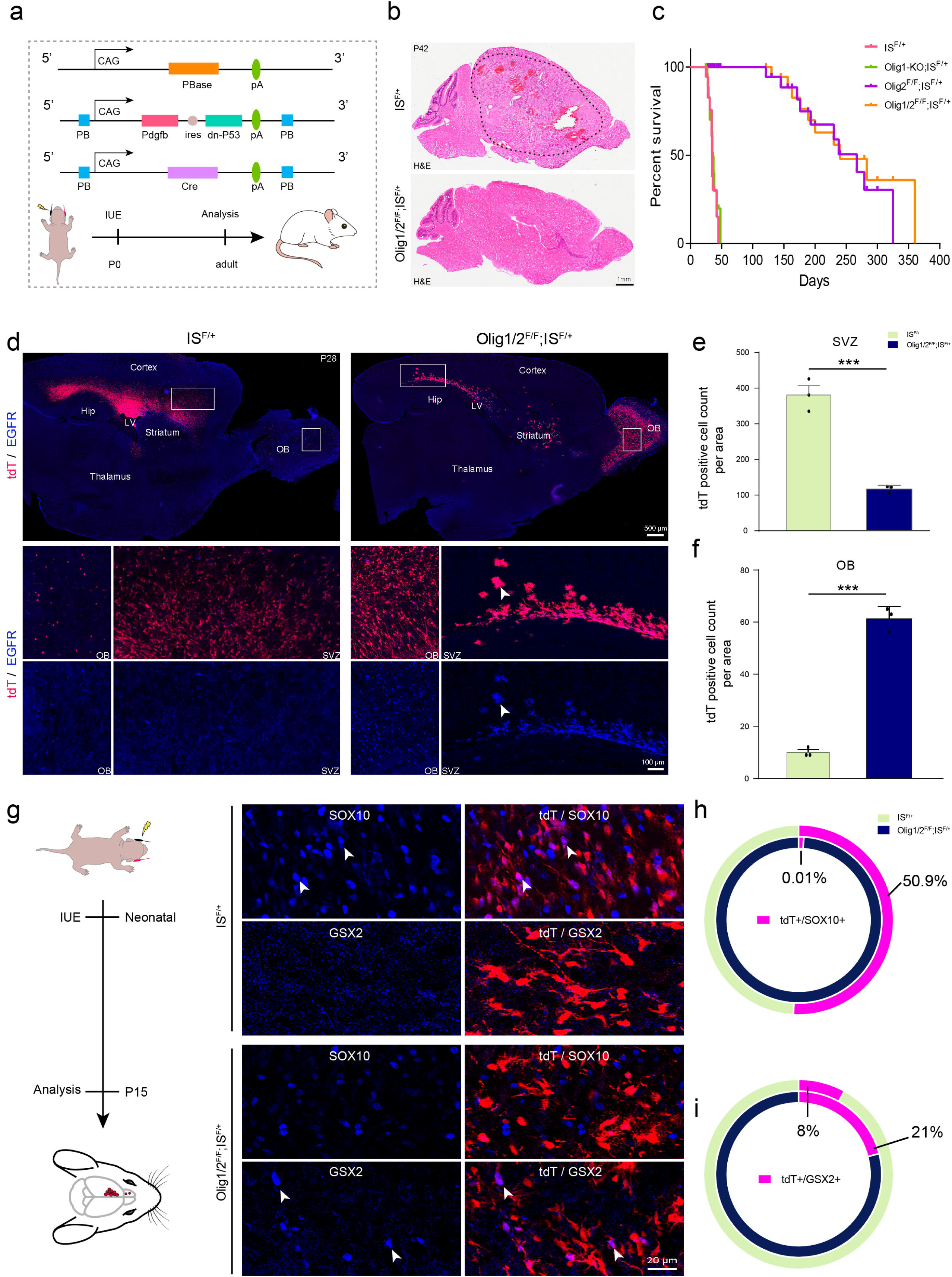
*Olig1/2* promote OL-lineage generation and inhibit the OB interneuron production through repressing *Gsx2* expression in glioblastoma. (a) A schematic diagram of construct mouse glioblastoma model. (b) Analysis of glioblastoma progression in *IS^F/+^* reporter mice and *Olig1/2^F/F^; IS^F/+^* mice by H&E staining at P42. (c) Kaplan-Meier survival analysis of *IS^F/+^* (n=12), *Olig1*-KO*; IS^F/+^* (n=12), *Olig2^F/F^; IS^F/+^* (n=10), and *Olig1/2^F/F^; IS^F/+^* (n=10) mice of the glioblastoma model from P10. (d) Immunofluorescence staining showing the distribution of the tdT- and EGFR-positive cells in the sagittal section of the *IS^F/+^* and *Olig1/2^F/F^; IS^F/+^* glioblastoma model mice. (e) The tdT- positive cell numbers are reduced in the dSVZ or cortex of the *Olig1/2^F/F^; IS^F/+^* glioblastoma model mice at P28, compared to *IS^F/+^* glioblastoma model mice. (f) The tdT- positive cell numbers are significantly increased in the OB of the *Olig1/2^F/F^; IS^F/+^* glioblastoma model mice at P28, compared to *IS^F/+^* glioblastoma model mice. (g) Immunofluorescence staining showing the tdT/GSX2 and tdT/SOX10 double-positive cells in the *IS^F/+^* and *Olig1/2^F/F^; IS^F/+^* glioblastoma model mice. (h) The ratio of tdT/SOX10 double-positive cells in and *Olig1/2^F/F^; IS^F/+^* and *IS^F/+^* glioblastoma model mice. (i) The ratio of tdT/GSX2 double-positive cells in *Olig1/2^F/F^; IS^F/+^* and *IS^F/+^* glioblastoma model mice. ***P<0.001 (one-way ANOVA followed by Tukey–Kramer post-hoc test). N=3 mice per group, mean±s.e.m.

Next, to investigate why the loss of *Olig2* or *Olig1/2* significantly extends the mouse survival time, we analyzed mice of different genotypes at P30. In our system, the tdTomato (tdT)^+^ cells are *Olig1*-ablated or *Olig2*-ablated or *Olig1/2*-ablated cells (Figure 5d and Figure S5a). As observed during normal development, we found that the number of the tdT-positive cells significantly increased in the OB of the *Olig2*-ablated and *Olig1/2*-ablated glioblastoma mice, compared to glioblastoma mice (Figures 5d-f and Figures S5a-c). We also found that the number of the tdT-positive cells increased in the OB of the *Olig1*-ablated glioblastoma mice, compared to glioblastoma mice (Figures 5d-f and Figures S5a-c). On the contrary, the number of the tdT-positive cells are greatly reduced in the SVZ (or cortex) of the *Olig2*-ablated and *Olig1/2*-ablated glioblastoma mice, compared to glioblastoma mice (Figures 5d-f and Figures S5a-c). A landmark study indicates that *Olig2* deletion in glioblastoma changes the gene expression profile from proneural (oligodendrocyte) to astrocyte-associated classical phenotype and leads to increased *Egfr* expression^62^. Indeed, we observed that *Egfr* expression was significantly upregulated in tdT-positive cells in the cortex and OB of *Olig2*-ablated and *Olig1/2*-ablated glioblastoma mice. However, these cells in the OB are OB neuron-like cells, not astrocyte-like glioblastoma cells, as these cells exhibit the morphology of mature OB interneurons (Figure 5d and Figure S5a).

To understand the molecular mechanisms behind this, we analyzed the glioblastoma model mice at P15, when the gliomas had just initiated^63^ (Figure 5g). Many tdT-positive cells co-expressed with OL-lineage marker SOX10, and fewer co-expressed with GSX2 (OB-lineage) in the *IS^F/+^* glioblastoma mice. In stark contrast, the number of tdT-positive cells co-expressing GSX2 are significantly increased, while those co-expressing SOX10 are significantly decreased in the *Olig1/2*-ablated glioblastoma mice (Figures 5g-i). Altogether, our findings suggest that *Olig1/2* deficiency causes tumor-initiating cells transit from producing highly proliferative OPC-like cells to producing proliferation-restricted OB interneurons.

### 6. Human H3.3G34R/V gliomas originate from cortical GPCs rather than from LGE progenitors

According to our above research, *Olig1/2* can promote OPC generation and inhibit OB interneuron generation in glioblastoma. A particular type of tumor that caught our attention is the H3.3-G34R/V pediatric high-grade gliomas, which is a newly recognized universally lethal childhood brain tumor^21^. Two studies indicate that H3.3-G34R/V gliomas originate from GSX2-positive progenitors in the LGE/MGE, offering important insights into their origins^38,42^. However, our recent studies showed that cortical-derived progenitors also expressed *Gsx2*^6,44^. Therefore, we hypothesize that cortical-derived GSX2-positive progenitors are the origin of H3.3-G34R/V gliomas.

To address this issue, we first analyzed the differentially expressed genes (DEGs) between H3.3-G34R/V and H3.3-K27M^21^ to determine the molecular features of the H3.3-G34R/V gliomas (Figure S6a). We found that, compared to H3.3-K27M, H3.3-G34R/V gliomas show high levels of *DLX2*, *PROKR2*, and *SP8* expression, while exhibiting low levels of *GPR17*, *OLIG1*, and *OLIG2* expression (Figure 6a). KEGG analysis of the DEGs in H3.3-G34R/V gliomas revealed that the downregulated genes are primarily involved in the regulation of glial cell differentiation, gliogenesis, and oligodendrocyte differentiation. Conversely, the upregulated genes are mainly associated with OB interneuron differentiation, neural fate commitment, and cell fate specification (Figure 6b). This was consistent with the phenotype observed *Olig1/2* mutant mice.

**Fig. 6.**
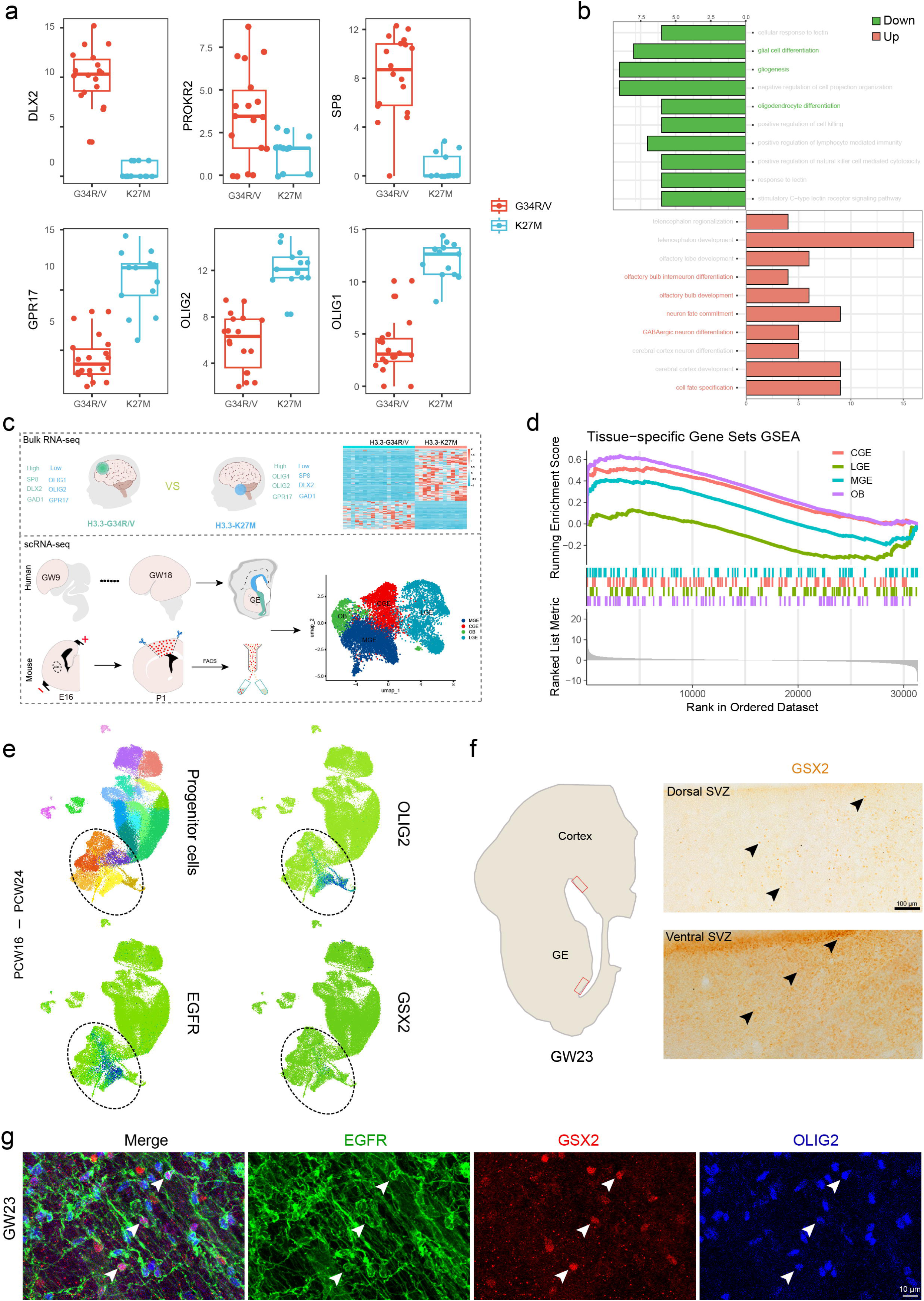
H3.3G34-mutant cells express high levels of genes relative to the OB-lineage and low levels of genes relative to the OL-lineage. (a) The expression of the *DLX2*, *PROKR2*, and *SP8* are increased, whereas the expression of the *GRP17*, *OLIG1*, and *OLIG2* are decreased in the H3.3G34R/V mutants, compared to H3.3K27M mutants. (b) The KEGG analysis of the DEGs between H3.3G34R/V mutants and H3.3K27M mutants. (c) Identifying differentially expressed genes (DEGs) between H3.3G34R/V mutants and H3.3K27M mutants (the upper part of C). Using the published human and mouse cortical scRNA-Seq data to identify the DEGs among the cortical-derived OB-lineage cells, LGE, MGE and CGE (the lower part of C). (d) GSEA analysis shows that DEGs in H3.3G34R/V mutants are more similar to those in cortical-derived OB-lineage cells than that in LGE, MGE and CGE. (e) The UMAP analysis shows that the human cortical-derived progenitor expressed *OLIG2*, *EGFR* and *GSX2* from PCW16 to PCW24. (f) *GSX2* is expressed in the ganglionic eminence and cortex of the human fetal brain at GW23. (g) The triple immunostaining of *EGFR*, *OLIG2* and *GSX2* in the human fetal brain at GW23.

Next, we performed integrative analysis of published human GE data^64^ and mouse cortex data^7^ to identify the molecular characteristics of LGE, MGE, CGE and cortical-derived OB-lineage cells (Figure 6c and Figures S6b-c). Using the GSEA and Upset analysis, we found that the molecular features of the H3.3-G34R/V gliomas are more inclined towards OB-lineage cells rather than the LGE, MGE and CGE (Figure 6d and Figure S6d). To further confirmed H3.3-G34R/V gliomas may originate from cortical-derived GSX2^+^ cells, we then reanalyzed the published human cortical scRNA-Seq data^60^ and found that cortical-derived progenitors express *PAX6*, *EOMES*, *NEUROG2*, *OLIG1*, *OLIG2*, and *EGFR* (Figure 6e and Figure S6e). A small population of cortical progenitors also express *GSX2* (Figure 6e). Indeed, immunohistochemical staining revealed high expression of *GSX2* in the cortex at GW23 (Figure 6f and Figure S6f). These GSX2-positive cells also express GPC markers, such as *EGFR* and *OLIG2* (Figure 6g). Our results indicated that cortical-derived progenitors can also express *GSX2* in humans. Moreover, PPI (protein-protein interaction) analysis of the DEGs identified key regulatory roles for *OLIG1/2*, *FOXG1*, and *DLX2* in H3.3-G34R/V gliomas (Figure S6g). These results indicated that due to lack of *OLIG1/2*, the H3.3-G34R/V gliomas may upregulate *GSX2* and generate more OB interneuron-like cells in the cortex. Taken together, our findings provide evidence that H3.3-G34R/V gliomas may originate from cortical GPCs rather than from LGE/MGE progenitors.

## DISCUSSION

Cortical gliogenesis is a complex process involving multiple signaling pathways and transcription factors that work together to regulate the generation of various cell types, including glial cells and neurons. The precise molecular mechanisms governing this process have yet to be fully elucidated. Our study reveals novel functions of *Olig1/2* in cortical gliogenesis that diverge from previous findings. We provide compelling evidence that *Olig1/2* have dual roles during cortical gliogenesis: they promote oligodendrocyte precursor cell (OPC) generation while repressing olfactory bulb (OB) interneuron production by directly inhibiting *Gsx2* expression through distinct enhancers in both mice and humans. Furthermore, we found that the absence of *Olig1/2* led to tumor-initiating cells ceasing the production of highly proliferative OPCs and instead transforming to generate proliferation-restricted olfactory bulb interneurons, suggesting the conserved roles of *Olig1/2* in both gliogenesis and glioblastoma. Additionally, our findings indicate that H3.3-G34R/V gliomas originate in cortical progenitors, rather than LGE/MGE progenitors. Therefore, our study not only revised the origin of H3.3-G34R/V gliomas but also emphasizes the need for a paradigm shift in how we approach glioblastoma treatment. By integrating insights from developmental biology with glioma research, we can begin to unravel the complexities of these tumors and identify more effective therapeutic targets.

The interactions between transcription factors are intricate and multifaceted. On the one hand, the homeobox gene *Gsx1/2*, for instance, control the timing of OPC specification within LGE progenitors by inhibiting *Olig2* expression^65,66^. In *Gsx1/2* double mutant mice, progenitors prematurely express *Olig2*, accelerating oligodendrocyte fate determination. On the other hand, *Gsx1/2* promote the expression of the *Dlx1/2* and then the *Dlx1/2* further inhibit the expression of *Olig1/2* during GABAergic interneurons development^57,59,67–71^. Previous studies have shown that *Olig1* directly represses *Dlx1/2* by regulating the I12b intergenic enhancer in the medial ganglionic eminence (MGE). However, *Olig2*-null animals exhibit no abnormalities in early interneuron development, likely due to the perinatal lethality of these mice, which limits the analysis of later developmental stages^72,73^. Actually, *Olig1* and *Olig2* have similar structures, it is very likely that they perform similar functions in OB interneurons, such as *Gsx1/2*, *Dlx1/2*, *Nr2f1/2*, and *Sp8/9* functions in OB neurons^58,65,74,75^. In this study, we found *Olig1/2* plays a dual role in transcriptional regulation in GPCs. On one hand, as a transcriptional activator, it activates the OPC production program, including genes like *Pdgfra*, *Sox8* and *Sox10*. On the other hand, as a transcriptional repressor, it inhibits the production of OB interneurons by suppressing *Gsx2* and *Dlx1/2*. What’s more, our study confirm for the first time that both *Olig1* and *Olig2* synergistically regulate the expression of *Gsx2* to control the fate commitment of cortical GPCs during gliogenesis.

Gliomas exhibit several early developmental characteristics, such as rapid cell proliferation, activation of developmental signaling pathways, significant cellular plasticity, and responsiveness to local environmental cues^61,76^. *Olig1/2* are ubiquitously expressed in gliomas and play critical roles in tumorigenesis and tumor phenotype plasticity^20,62,77^. Recent studies have shown that *Olig1/2*-expressing intermediate lineage progenitors are predisposed to PTEN/p53-loss–induced gliomagenesis and harbor specific therapeutic vulnerabilities^19^. *Olig2*, in particular, plays extensive and pivotal roles across various glioma subtypes^35,62^. Interestingly, one study reported *Olig2* has the capacity to dictate glioma subtype, as a loss of *Olig2* leads to a shift from a proneural transcriptional subtype towards a more astrocytic phenotype, including downregulation of *Pdgfra* and concomitant upregulation of *Egfr*. Another study showed that abrogation of *Olig2* function in glioma stem-like cells (GSCs) cause a shift from proneural-to-mesenchymal gene expression pattern^35^. However, in our *Olig1/2*-CKO glioma mouse model, we observed that tumor formation is disrupted, resulting in the generation of many OB neurons, despite the high expression levels of *Egfr* in these tumor-like cells (Figure5 and FigureS5). Thus, the role of *Olig1/2* in tumorigenesis is context-dependent, possibly involving the inhibition of *Egfr* expression in different glioma subtypes. It is worth noting that the functions of OLIG1/2 are also closely associated with their post-transcriptional regulatory mechanisms. For example, phosphorylation of a highly conserved triple serine motif at the N-terminus of OLIG2 (S10, S13, and S14) is crucial for regulating glioma cell proliferation and invasion.^78–80^.

Identifying the developmental origins of gliomas can greatly help our understanding of tumor biology. Pioneering research has indicated that H3.3G34R/V gliomas originate from GSX2-positive progenitors located in the LGE with PDGFR mutations^42^. In fact, these gliomas are typically found in the cerebral cortex rather than the basal ganglia^40,81^. Furthermore, our recent studies have shown that *Gsx2* is also expressed in cortical-derived GPCs during gliogenesis^6,44^. In this study, we found that *GSX2* is expressed in human cortical GPCs, and the gene expression profile of H3.3G34R/V gliomas is similar to that of cortical-derived OB neurons (Figure6 and FigureS6). Given the lack of *OLIG1/2* expression and enrichment of *GSX2-DLX1/2-PROKR2* expression in H3.3G34R/V gliomas, we speculate that these gliomas originate from cortical GSX2-positive GPCs.

In summary, our findings uncovered the molecular mechanisms by which the transcription factors *Olig1/2* determine cell fate in gliogenesis and gliomagenesis. These findings provide new insights into potential therapies for gliomas, suggesting that interfering with the expression of *Olig1/2* could transform proliferative OPC-like tumor cells into non-proliferative OB interneuron-like cells. Moreover, we also provide new evidence that the H3.3G34R/V glioma are derived from the cortical-derived GPCs. These findings provide a bridge for future research to understanding the relationship between normal development and gliomagenesis.

## MATERIALS AND METHODS

### KEY RESOURCES TABLE

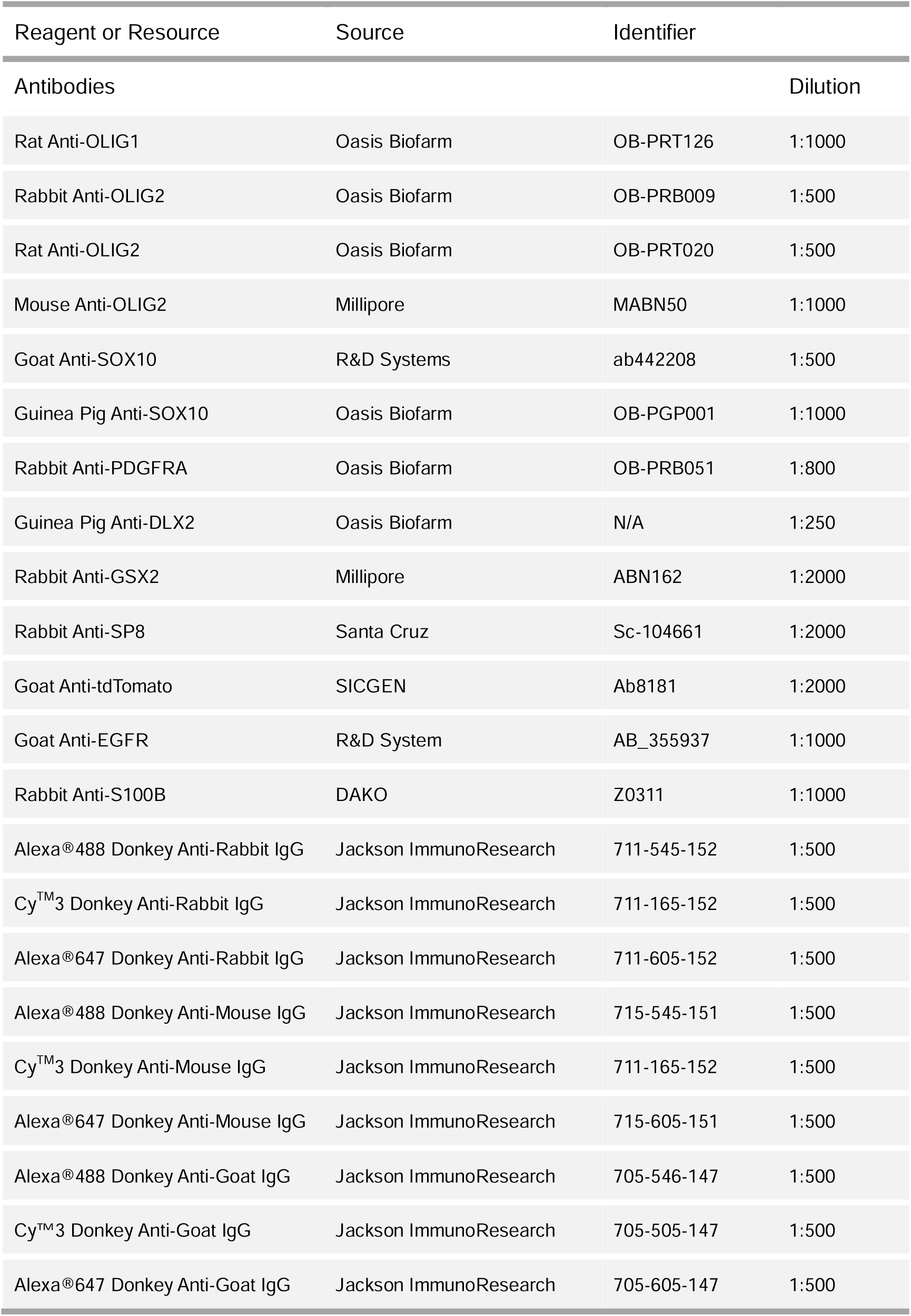
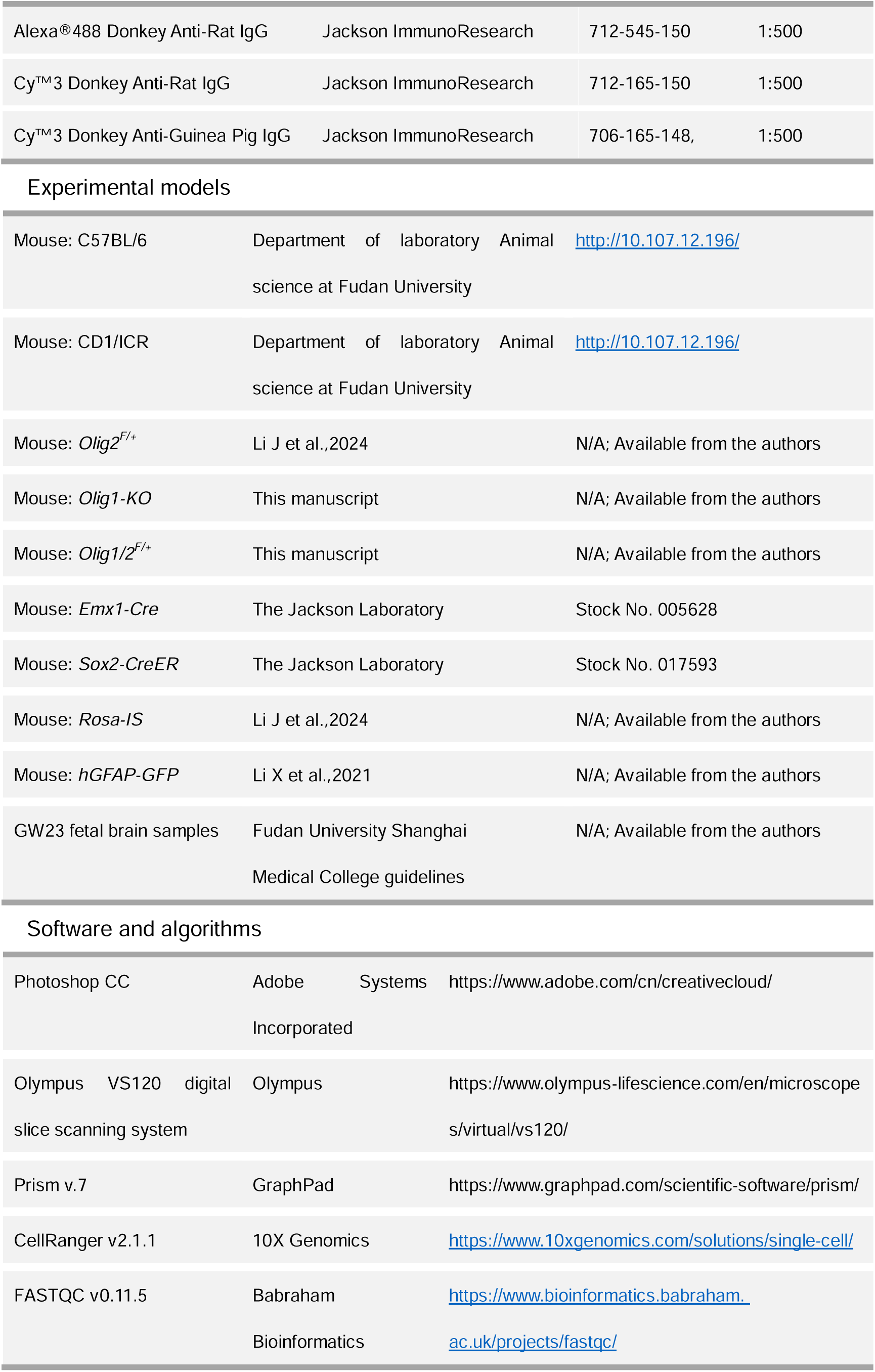
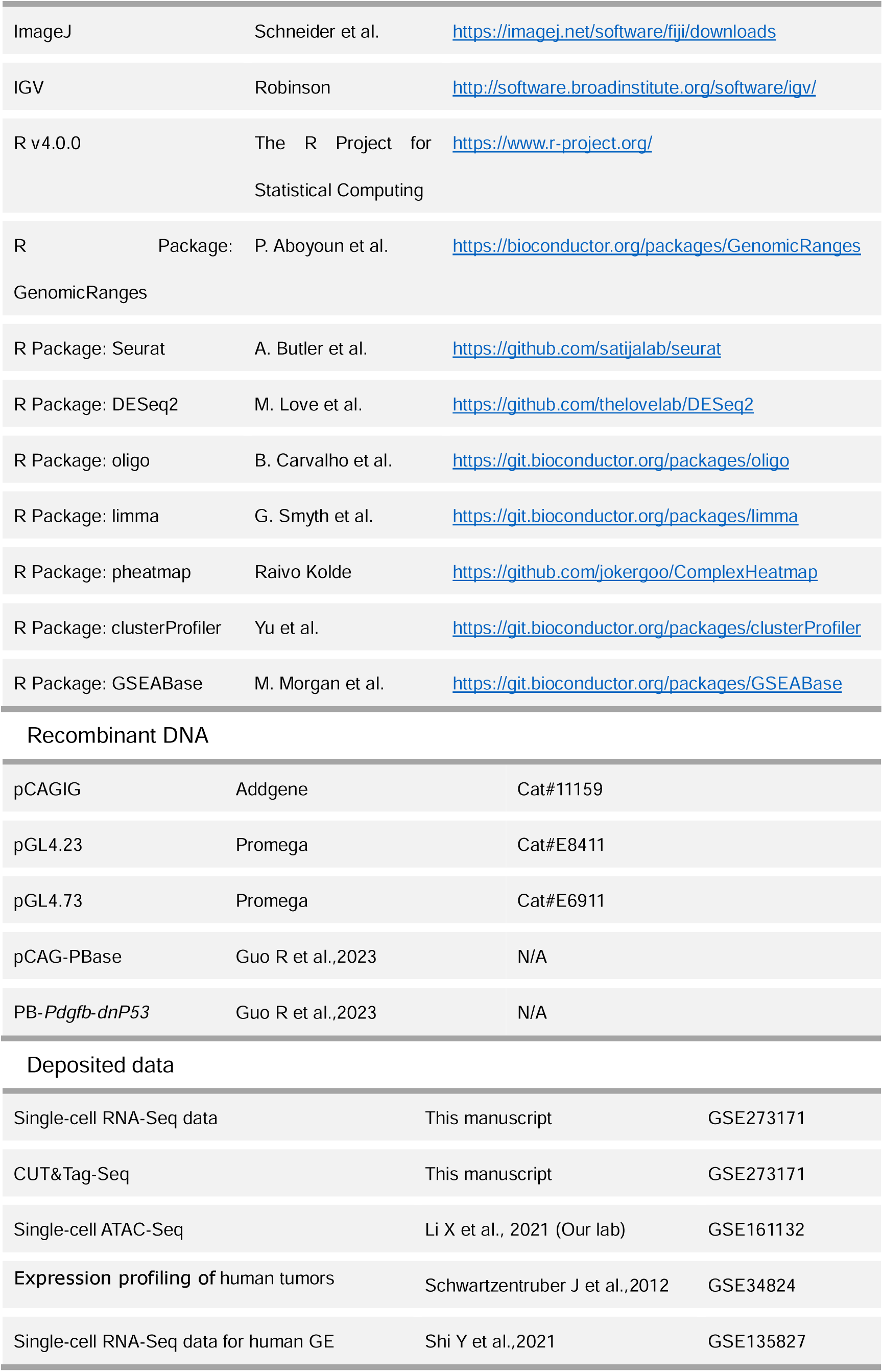
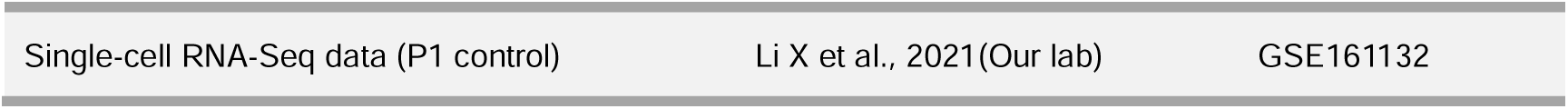

### RESOURCE AVALABILITY

#### Lead Contact

Further information and requests for resources and reagents should be directed to and will be fulfilled by the Lead Contact, Zhuangzhi Zhang.

#### Material Availability

This study did not generate new unique reagents.

#### Data and Code Availability

The accession numbers for the RNA-Seq and CUT&Tag-Seq datasets reported in this paper are Gene Expression Omnibus (GEO): GSE273171

### EXPERIMENTAL MODEL AND METHOD DETAILS

#### Animals

All experiments conducted in this study were in accordance with guidelines from Fudan University (No. 20220228- 140). *Olig2^F/+^*^44^, *Rosa-IS*^44^, *hGFAP-GFP*^44^, *Sox2-CreER*^82^, *Emx1-Cre*^83^ mice were previously described. *Olig1/2^F/+^* mice were generated via the CRISPR/Cas9 strategy. Lox2272 sites flanked the coding region of two exons of *Olig2* and LoxP sites flanked the one exon of *Olig1*. *Olig1*-KO mice were generated via the CRISPR/Cas9 strategy. LoxP sites flanked the one exon of OLIG1 (Figures S1a). Wild-type mice or littermates without the *Cre* allele were used as controls. All mice were maintained on a mixed genetic background of C57BL/6J and CD1. The mice were allowed access to water and food ad libitum and maintained on a 12-hour light/dark cycle. The day of vaginal plug detection was considered embryonic day 0.5 (E0.5), and the day of birth was defined as postnatal day 0 (P0). Both male and female mice were used in all experiments.

#### Tamoxifen Labeling

A single intraperitoneal injection of Tamoxifen (100 mg/kg) was administered at P12 or P14 and mouse was analyzed 7 days after TM injection.

#### Tissue Preparation

Mice were deeply anesthetized and perfused intracardially thoroughly with ice-cold PBS followed by 4% paraformaldehyde (PFA). All brains were fixed overnight in 4% PFA at 4°C, cryoprotected in 30% sucrose for at least 24 hours, embedded in optimal cutting temperature (O.C.T.) (Sakura Finetek) on dry ice and ethanol slush and preserved at -80 °C. Human tissue preparation was performed as previously described^8^.

#### Immunofluorescence

In this study, 20-μm thick frozen sections were used for immunostaining. The sections were washed with 0.05 M TBS for 10 min, incubated in Triton-X-100 (0.5% in 0.05 M TBS) for 30 min at room temperature (RT), and then incubated with blocking solution (10% donkey serum +0.5%; Triton-X-100 in 0.05 M TBS, pH= 7.2) for 2 h at RT. The primary antibodies were diluted in 10% donkey serum blocking solution, incubated overnight at 4 °C, and then rinsed 3 times with 0.05 M TBS. Secondary antibodies (from Jackson, 1:500) matching the appropriate species were added and incubated for 2-4 h at RT. Fluorescently stained sections were then washed 3 times with 0.05 M TBS. This was followed by 4’,6-diamidino-2-phenylindole (DAPI) (Sigma, 200 ng/ml) staining for 5 min, and then the sections were then cover-slipped with Gel/Mount (Biomeda, Foster City, CA).

#### In Situ Hybridization

ISH was performed as previously described^84^. Briefly, ISH was performed on 20-μm cryosections using digoxigenin riboprobes. Probes were made from P0 wild-type mouse brain cDNA amplified by PCR using the following primers:

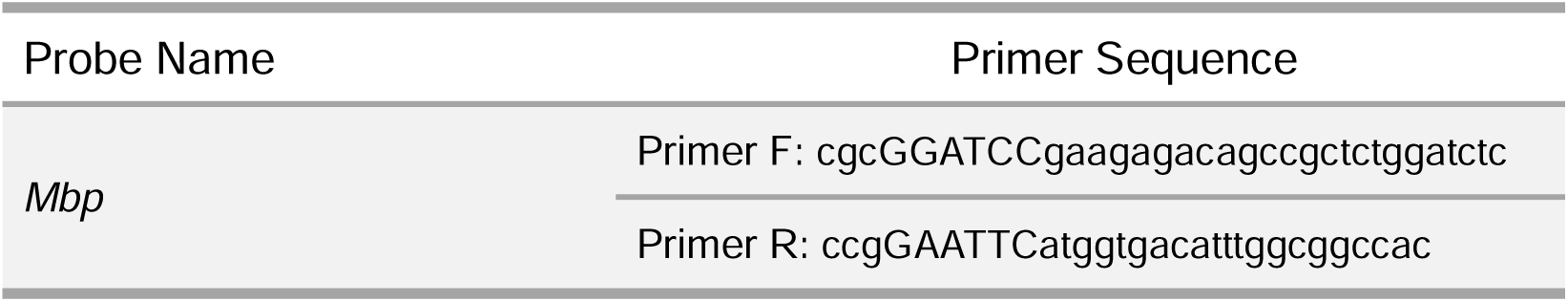

#### In utero electroporation (IUE)

In utero electroporation (IUE) of IS embryos was performed at E16.5 or P0. The pCAG-Cre, pCAG-PBase or PB-Pdgfb-dnP53 plasmid was mixed with 0.05% Fast Green (Sigma) and injected into the lateral ventricle of embryos (0.5 μL per embryo) using a beveled glass micropipette. Five electrical pulses (duration: 50 ms) were applied at 42 V across the uterine wall with a 950 ms interval between pulses. Electroporation was performed using a pair of 5 mm platinum electrodes (BTX, Tweezertrode 45-0488, Harvard Apparatus) connected to an electroporator (BTX, ECM830).

#### Plasmid Construction

The pCAG-*Cre* vector was constructed as reported. The coding sequence of *Cre* in the pCAG-*Cre* vector was replaced by the PiggyBac transposase to construct the pCAG-PiggyBac transposase vector. Internal ribosomal entry site sequence was placed between the targeted CDS and the GFP reporter. For tumor-inducing vectors, *Pdgfb* was cloned into the downstream of the CAG promoter, and GFP was replaced by an HA-tagged dn*P53*.

#### scRNA-seq

After anesthetizing the mice, make a T-shaped incision on the back of the head. Use forceps to peel back the skin and skull on both sides to expose the brain tissue. Insert the forceps into the base of the brain and remove the brain, placing it in HBSS. If needed, clip the tail for PCR identification. Under a dissecting microscope, peel off the meninges, cut the target tissue (cortex) into small pieces. Transfer the tissue pieces to a 15 ml centrifuge tube using a 1 ml pipette. After standing for 30 seconds, remove the excess HBSS. Add 2 ml of preheated (37°C) cell dissociation solution to the tissue pieces. Incubate in a 37°C incubator for 20 minutes, gently shaking every 5 minutes to mix. Add 2 ml of 10% FBS-DMEM to stop the reaction. Pipette up and down (10-15 times) until a single-cell suspension is formed. Rinse a 40 μm MACS SmartStrainer (filter) with 10% FBS-DMEM. Place the filter on a 15 ml centrifuge tube and filter the cell suspension through it. Centrifuge at 500 g for 5 minutes at 4°C. Remove the supernatant and resuspend the cells in an appropriate amount of 10% FBS-DMEM. Keep on ice until use. According to the experimental design, perform flow cytometry sorting on the concentrated cells to obtain positive target cells. After obtaining single-cell suspensions from the target mice (*Olig1/2^f/f^; IS^F/+^*), we used the Countstar instrument to count the cells and assess viability (trypan blue staining), ensuring the quality and viability of the cells. Once qualified, the library construction and sequencing were outsourced to an external company. The single-cell sequencing library was constructed using 10X Genomics technology and high-throughput sequencing was performed on the Illumina Hiseq4000. After sequencing, we used CellRanger for quality control and preprocessing of the raw data to ensure the reliability of subsequent analysis. Through this series of steps, we obtained the cell expression matrix file, which was then used for single-cell bioinformatics analysis using the Seurat package in R.

#### CUT&Tag-Seq

The CUT&Tag assay was conducted on FACS-sorted GFP^+^ cells, using the cortex dissected from *hGFAP*-GFP at perinatal stages. Initially, 1×10^5 cells were washed with 500 μl of wash buffer (10× wash buffer (-), 50× protease inhibitor cocktail, ddH2O) and centrifuged at 600 rcf for 3 minutes at room temperature (RT). The cell pellets were then resuspended in 100 μl of wash buffer. Concanavalin A-coated magnetic beads were washed twice with binding buffer (10× binding buffer, ddH2O). Next, 10 μl of activated beads were added and incubated at RT for 10 minutes. Bead-bound cells were resuspended in 50 μl of antibody buffer (10× wash buffer (-), 50× protease inhibitor cocktail, ddH2O, 5% digitonin, 0.5 M EDTA, 30% BSA). Then, 1 μg of primary antibody (mouse polyclonal anti-Olig2 and anti-Olig1) or no antibody (control) was added and incubated at RT for 2 hours. The primary antibody was removed using a magnetic stand. The secondary antibody was diluted in 50 μl of Dig-wash buffer (10× wash buffer (-), 50× protease inhibitor cocktail, ddH2O, 5% digitonin), and the cells were incubated at RT for 1 hour. Cells were washed three times with Dig-wash buffer to remove unbound antibodies.

The Hyperactive pG-Tn5 Transposase adaptor complex (TTE mix, 4 μM, Vazyme) was diluted 1:100 in 100 μl of Dig-300 buffer (10× Dig-300 buffer (-), 5% digitonin, 50× protease inhibitor cocktail, ddH2O). The cells were incubated with 0.04 μM TTE mix at RT for 1 hour and then washed three times with Dig-300 buffer to remove unbound TTE mix. The cells were resuspended in 300 μl of tagmentation buffer (10 mM MgCl2 in Dig-300 buffer) and incubated at 37 °C for 1 hour. To terminate tagmentation, 10 μl of 0.5 M EDTA, 3 μl of 10% SDS, and 2.5 μl of 20 mg/ml Proteinase K were added to the 300 μl sample and incubated overnight at 37 °C. DNA was purified using phenol-chloroform-isoamyl alcohol extraction and ethanol precipitation with RNase A treatment.

For library amplification, 24 μl of DNA was mixed with 1 μl of TruePrep Amplify Enzyme (TAE, Vazyme), 10 μl of 5× TruePrep Amplify Enzyme Buffer, 5 μl of ddH2O, and 5 μl of uniquely barcoded i5 and i7 primers from the TruePrep Index Kit V2 for Illumina (Vazyme). A total of 50 μl of the sample was placed in a thermocycler. To purify the PCR products, 1.2× volumes of VAHTS DNA Clean Beads (Vazyme) were added and incubated at RT for 10 minutes. The libraries were washed twice with 80% ethanol and eluted in 22 μl of ddH2O. The libraries were sequenced on an Illumina NovaSeq platform, producing 150-bp paired-end reads. All raw sequence data were quality trimmed to a minimum Phred score of 20 using Trimmomatic. Apparent PCR duplicates were removed using Picard MarkDuplicates vl.107. All reads generated from the CUT&Tag-Seq of OLIG1 and OLIG2 were aligned to the mm39 mouse genome using Bowtie2 version 2.3.4. Sequence tags were aligned to the genome and subsequently analyzed by MACS2 software version 2.1.4 to detect genomic regions enriched for multiple overlapping DNA fragments (peaks), which were considered putative binding sites. Peaks with a false discovery rate lower than 5% were retained for further chromosomal region analysis. Visualization of peak distribution along genomic regions of genes of interest was performed using the Integrative Genomics Viewer (IGV).

#### Dual-luciferase Assay

The DNA fragments of the *Gsx2*-DSG-1, *Gsx2*-DSG-2, *Gsx2*-DSG-3, *Gsx2*-hs687, and *Gsx2*-PE near the *Gsx2* gene were created by PCR and subsequently cloned into the pGL4.23 firefly luciferase vector (Promega) upstream (U) or downstream (D) of the Luc2 gene. Cells from the mouse embryonal carcinoma cell line 293T were grown in DMEM (Gibco, 12571063) supplemented with 10% fetal bovine serum (FBS) (Gibco, 10099141). For the luciferase assay, 293T cell transfections were performed in triplicate in 24-well plates by using Lipo8000 transfection reagent according to the manufacturer’s protocol (Beyotime, C0533). Luciferase activity was quantified by a microplate luminometer (Turner BioSystems, Modulus microplate reader). The primers used for amplifying the putative *Gsx2* enhancers were as follows:

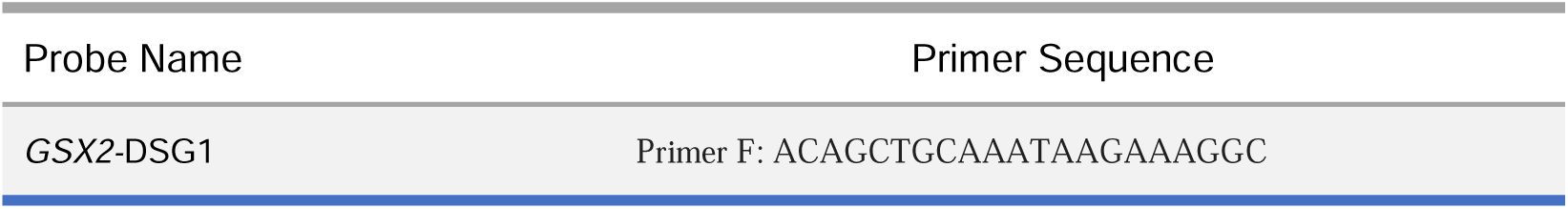
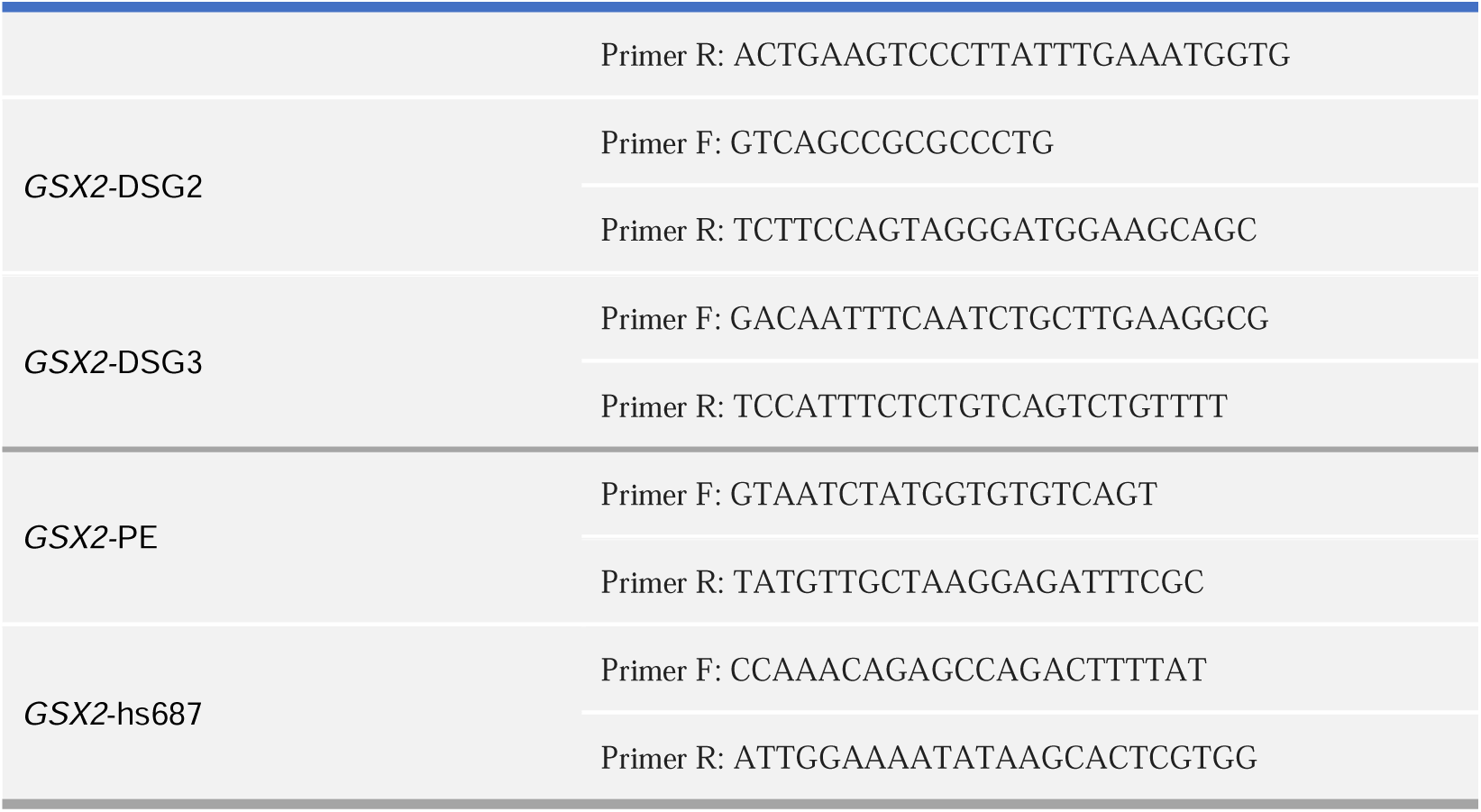

#### Image Acquisition and Statistical Analysis

Stained tissue sections were imaged on an Olympus VS 120 microscope or an Olympus FV3000 confocal microscope system. Images were processed by Adobe Photoshop CC and ImageJ software without distorting the original information.

The numbers of SP8^+^ and DLX2^+^ and OLIG2^+^/SOX10^+^ cells in the dorsal SVZ (1000×500 pixels area for the cortex imaged on Olympus FV3000), were quantified in 3-4 randomly chosen 20 mm sections for control (WT), *Olig2*-ECKO, *Olig1/2*-ECKO, *Olig1*-KO mice at P0,P7,P14 and P30.The numbers of SP8+ cells in the GL and GCL of OB(500 × 1000 pixels area imaged on Olympus VS 120) were quantified in 3-4 randomly chosen 20 mm sections for control and *Olig1/2*-ECKO at P7.The numbers of Mbp-positive cells in the corpus callosum and cortex (1000×1000 pixels area imaged × on Olympus VS 120) were quantified in 3-4 randomly chosen 20 mm sections for WT, *Olig2*-ECKO, *Olig1/2*-ECKO, *Olig1*-KO mice at P7 and P30. The numbers of tdT^+^/SOX10^+^, tdT^+^/PDGFRA^+^ and tdT^+^/SP8^+^ cells in the cortex (1000×500 pixels area imaged on Olympus FV3000) were quantified in 3-4 randomly chosen 20 mm sections for *IS^F/+^*, *Olig1*-KO; *IS^F/+^*, *Olig2^F/F^; IS^F/+^*, and *Olig1/2^F/F^; IS^F/+^* (e16@pCAG-*Cre*) mice at P7. The numbers of tdT^+^/EGFR^+^ and tdT^+^/SOX10^+^ in dorsal-SVZ and OB (500 × 500 pixels area for both areas imaged on Olympus FV3000) were quantified in 3-4 randomly chosen 20 mm sections in the sagittal section of *IS^F/+^, Olig1/2^F/F^; IS^F/+^*, *Olig1*-KO; *IS^F/+^, Olig1/2^F/F^; IS^F/+^* glioblastoma model mice at P15 and P28.

Analyses were performed using GraphPad Prism 6.0, Microsoft Excel and R language. For individual experiments, at least three samples of control or mutant mice were examined. Multiple sections from individual brains at similar positions were analyzed in morphological experiments. The analyzed mice were littermates of both sexes and age-matched. Values and error bars represent the mean±SEM. The respective replicate number (n) is indicated in the figures. P values were determined by an appropriate statistical test, such as a *student’s* t test for normally distributed data. P < 0.05 was defined as statistically significant. *P < 0.05, **P < 0.01, and ***P< 0.001.

## ACKNOWLEDGMENTS

This study was supported by the Ministry of Science and Technology of China (STI2030- 2021ZD0202300), National Natural Science Foundation of China (NSFC 31820103006, 32070971, 32200792, and 32200776), China Postdoctoral Science Foundation (2024T170179).

## AUTHOR CONTRIBUTIONS

Y.T. and Z.W. performed all experiments and analyzed data. F.Y., W.Z., J.L., Y.L., T.F., W.Z., Z.X., Y.Y., and X.S. helped to conduct experiments. J.S., Y.X., and Z.Y., help analyze the data. Z.Z. designed the experiments and analyzed data. Z.Z. and Z.Y. wrote the manuscript.

## Figure legends

**Fig. S1 *Olig1* and *Olig2* co-regulate the production of OB interneurons.**

(a) Schematic diagram of generating *Olig1* conventional knockout mice and *Olig2* and *Olig1/2* conditional knockout mice.
(b) The immunostaining of the SP8 (cyan) and DLX2 (red) in the dorsal SVZ of the control, *Olig1*-KO, *Olig2*-ECKO and *Olig1/2*-ECKO mice at P0.
(c) The sagittal section showing the DLX2 expression in the SVZ, RMS and OB of the control, *Olig1*-KO, *Olig2*-ECKO and *Olig1/2*-ECKO mice.
(d) Quantifications of the SP8-positive cells in OB GL and GCL of the control and *Olig1/2*-ECKO mice at P7. *P<0.05, **P<0.01 (one-way ANOVA followed by Tukey–Kramer post-hoc test, mean±s.e.m). N=4 mice per group.
(e) The immunostaining of the SP8 (cyan) and DLX2 (red) in the dorsal SVZ of the control and *Olig1/2*-ECKO mice at P30. OB, olfactory bulb; RMS, rostral migratory stream; SVZ, subventricular zone.

**Fig. S2 *Olig1/2* promote the production of cortical OPCs.**

(a) The triple immunostaining of the SOX10, OLIG2 and S100B in the control, *Olig1*-KO, *Olig2*-ECKO and *Olig1/2*-ECKO mice at P0.
(b) The majority of the SOX10/OLIG2 double positive cell are also expressed S100B in the *Olig2*-ECKO and *Olig1*/2-ECKO mice at P0, compared to the control and *Olig1*-KO mice.
(c) The expression pattern diagram of the *Sox10*, *S100b* and *Mbp* in the OL-lineage cells.
(d) The cluster 2 is the OL-lineage cells, which expressed high level *Sox10* at P7.
(e) The cluster 2 is expressed higher level of *S100b* in the *Olig1*/*2*-ECKO mice than that in the control mice at P7.
(f) The *Emx1* is expressed lower level in the *Olig1/2*-ECKO mice than that in the control mice at P7.
(g) The statistical data of the *Mbp* positive cells in the control, *Olig1*-KO, *Olig2*-ECKO and *Olig1/2*-ECKO mice both in corpus collosum and cortex at P30.
(h) *In situ* hybridization shows that *Mbp* expression is partial recovery of the *Olig1*-KO, *Olig2*-ECKO and *Olig1/2*-ECKO mice at P30, compared to control mice. ***P<0.001 (one-way ANOVA followed by Tukey–Kramer post-hoc test). N≥4 mice per group, mean±s.e.m. CC, corpus callosum.

**Fig. S3 *Olig1/2* promote OL-linage fate acquisition and repress OB-lineage fate acquisition in GPCs.**

(a) Both *Olig1* and *Olig2* are significantly reduced in the *Olig1/2^F/F^; IS^F/+^* (*Olig1/2*-CKO), compared to the *IS^F/+^* reporter (control) mice.
(b) The OL-lineage cells, which express *Pdgfra* and *Cntn1* are significantly decreased in the *Olig1/2*-CKO mice, compared to control mice.
(c) The OB-lineage cells, which express *Gad2* and *Sp9* are significantly increased in the *Olig1/2*-CKO mice, compared to control mice.
(d) The GPCs, which express *Egfr* and *Ascl1* are significantly increased in the *Olig1/2*-CKO mice, compared to control mice.
(e) Combining in utero electroporation (IUE) with immunohistochemistry to show tdT/SP8 double-positive cells in the dSVZ of the *IS^F/+^*, *Olig1*-KO; *IS^F/+^*, *Olig2^F/F^; IS^F/+^*, and *Olig1/2^F/F^; IS^F/+^* mice.
(f) The statistical data shows that the percentage of the tdT/SP8 double-positive cells are increased in the *Olig1*-KO; *IS^F/+^*, *Olig2^F/F^; IS^F/+^* and *Olig1/2^F/F^; IS^F/+^* mice, compared to control mice. *P<0.05, **P<0.01 (one-way ANOVA followed by Tukey–Kramer post-hoc test). N=3 mice per group, mean±s.e.m.

**Fig. S4 *Olig1/2* promote the generation of the cortical OL-lineage by regulating *Sox10* and *Pdgfra* expression.**

(a) Heatmap showing OLIG1 and OLIG2 binding sites in the promoter (<3 kb).
(b) The peak plot shows enrichment of OLIG1 and OLIG2 binding sites in the promoter region.
(c) The top 10 enriched known motifs of the OLIG1 and OLIG2 binding sites.
(d) Joint analysis of CUT&Tag-Seq and scATAC-Seq suggests that OLIG1 and OLIG2 cooperatively regulate the expression of *Pdgfra* by directly binding its promoter and potential enhancer.
(e) Combining CUT&Tag-Seq with scATAC-Seq to show that OLIG1 and OLIG2 cooperatively regulate the expression of *Sox8* in the cortex.
(f) Joint analysis of CUT&Tag-Seq and scATAC-Seq shows that OLIG1 and OLIG2 cooperatively regulate the expression of *Sox10* through directly binding multiple *Sox10* enhancers.
(g) The violin plot shows *Gsx2* expression is greatly increased in the GPCs of the *Olig1/2*-CKO mice, compared to control mice.
(h) Joint analysis of CUT&Tag-Seq and scATAC-Seq suggests that OLIG1 and OLIG2 cooperatively regulate the expression of *Gsx2* in GPCs by binding to potential enhancer (PE) near the *Gsx2* gene.

**Fig. S5 *Olig1/2* promote OL-lineage production and inhibit the OB interneuron production in glioblastoma.**

(a) Immunofluorescence staining showing the distribution of the tdT- and EGFR-positive cells in the sagittal section of the *Olig1*-KO; *IS^F/+^* and *Olig2^F/F^; IS^F/+^* glioblastoma model mice.
(b) The tdT- positive cell numbers are reduced in the dSVZ or cortex of the *Olig2^F/F^; IS^F/+^* glioblastoma model mice at P28, compared to *Olig1*-KO; *IS^F/+^* glioblastoma model mice.
(c) The tdT- positive cell numbers are significantly increased in the OB of the *Olig2^F/F^; IS^F/+^* glioblastoma model mice at P28, compared to *Olig1*-KO; *IS^F/+^* glioblastoma model mice. ***P<0.001 (one-way ANOVA followed by Tukey–Kramer post-hoc test). N=3 mice per group, mean±s.e.m.

**Fig. S6 H3.3G34-mutant gliomas likely originate from the cortex.**

(a) The heatmap showing the DEGs between H3.3G34R/V mutants and H3.3K27M mutants.
(b) UMAP analysis showing human ganglionic eminence (GE) cell clusters.
(c) The human MGE expressed *LHX6.* The LGE expressed *ISL1* and *SIX3*, and CGE expressed *SCGN* and *NR2F2*.
(d) The UpSet analysis shows that DEGs between H3.3G34R/V mutants and H3.3K27M mutants shared more genes with OB-lineage than that in LEG, MGE and CGE.
(e) The human cortical progenitor cells express *PAX6*, *EOMES*, *NEUROG2* and *OLIG1*.
(f) Immunohistochemistry for *GSX2* on GW23 brain section.
(g) The Protein-protein interaction (PPI) analysis showing that the OLIG1/2 play important roles in H3.3G34R/V mutants. LGE, lateral ganglionic eminence; MGE, medial ganglionic eminence; CGE, caudal ganglionic eminence.

