## Supplementary figures and images for "Orchestrating the Acquisition of Oligodendrocyte Precursor Cell versus Olfactory Bulb Interneuron Fates through *Olig1/2* during Mammalian Cortical Gliogenesis and Gliomagenesis"

### Supplemental Figure1

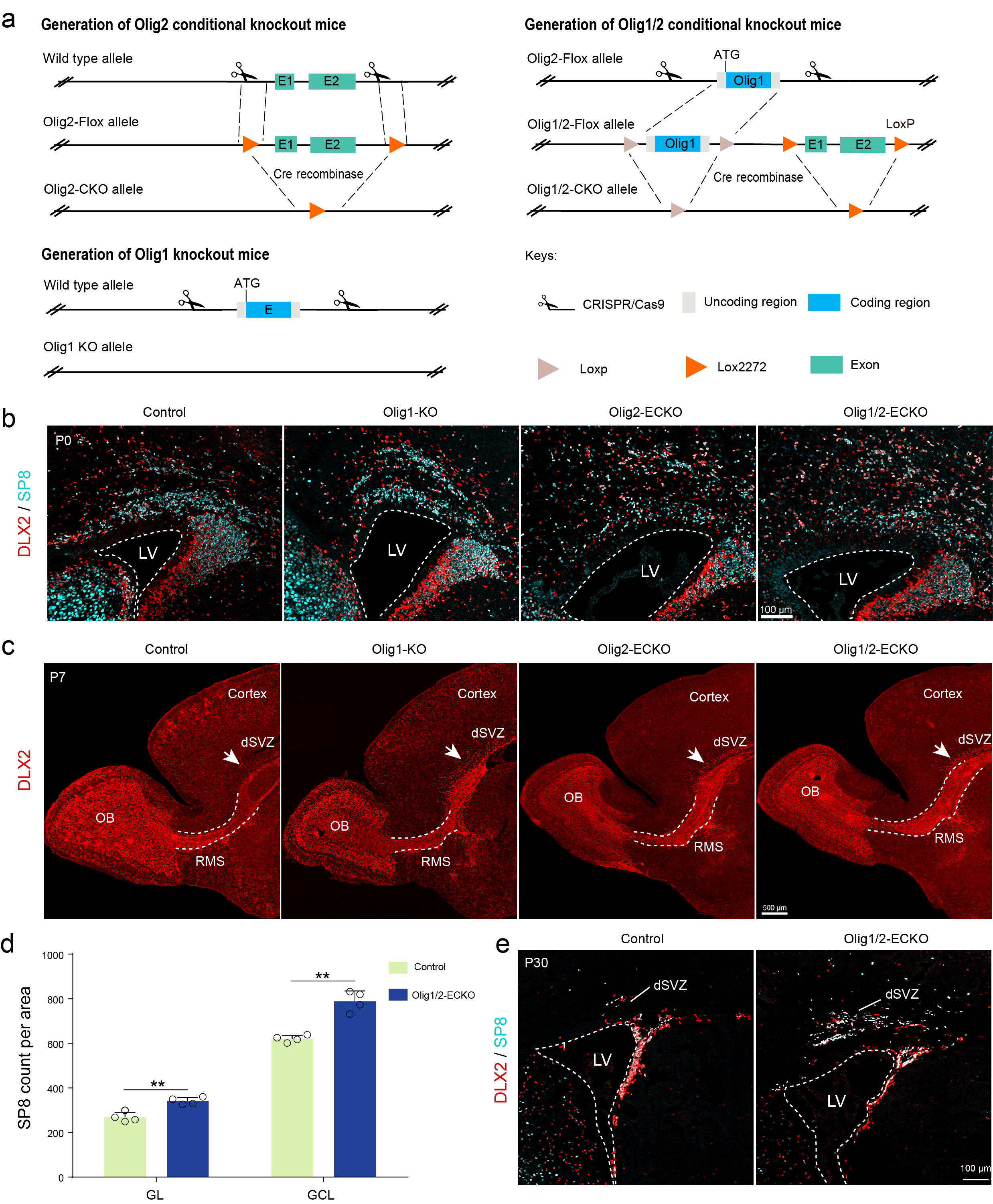

### Supplemental Figure2

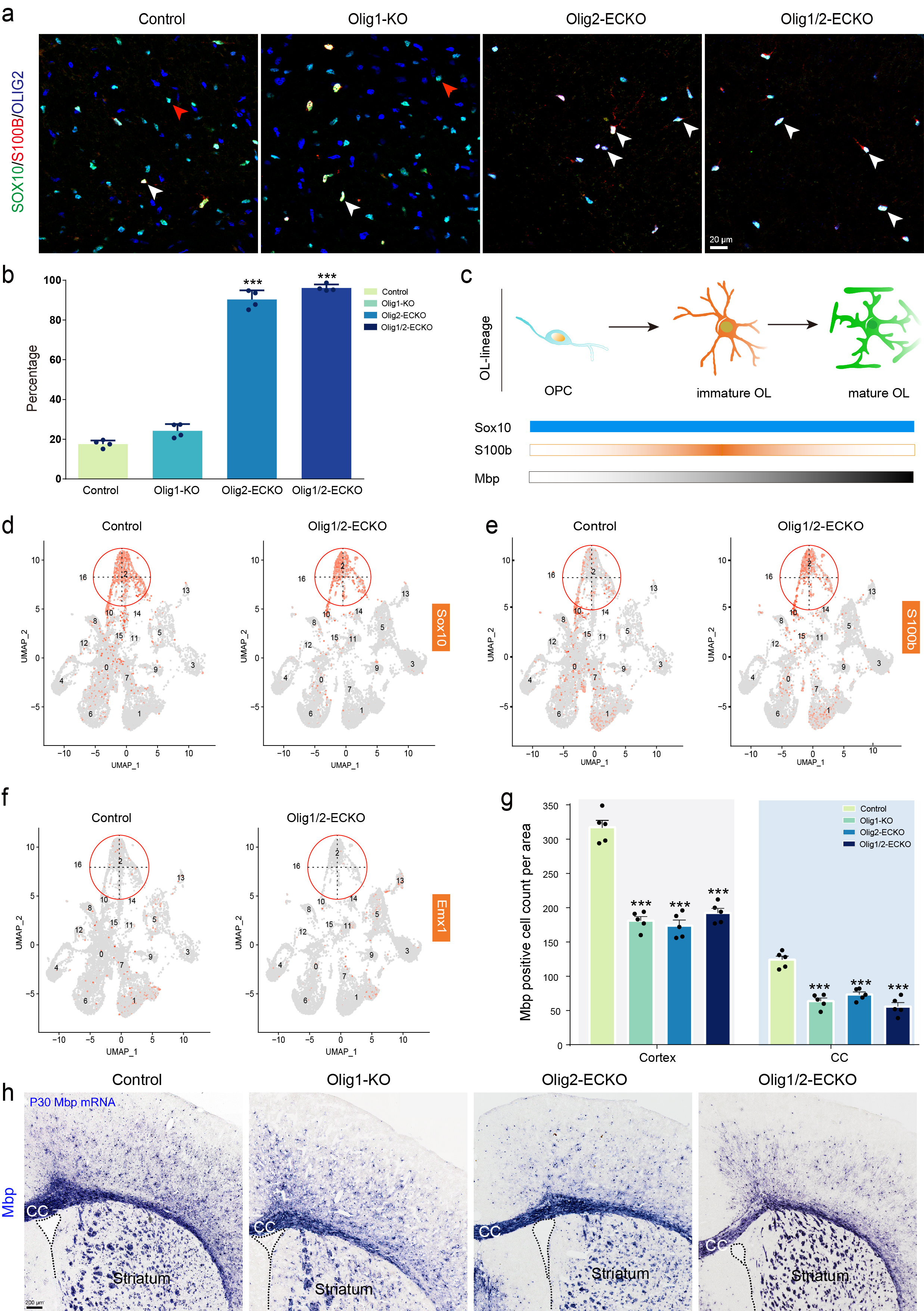

### Supplemental Figure3

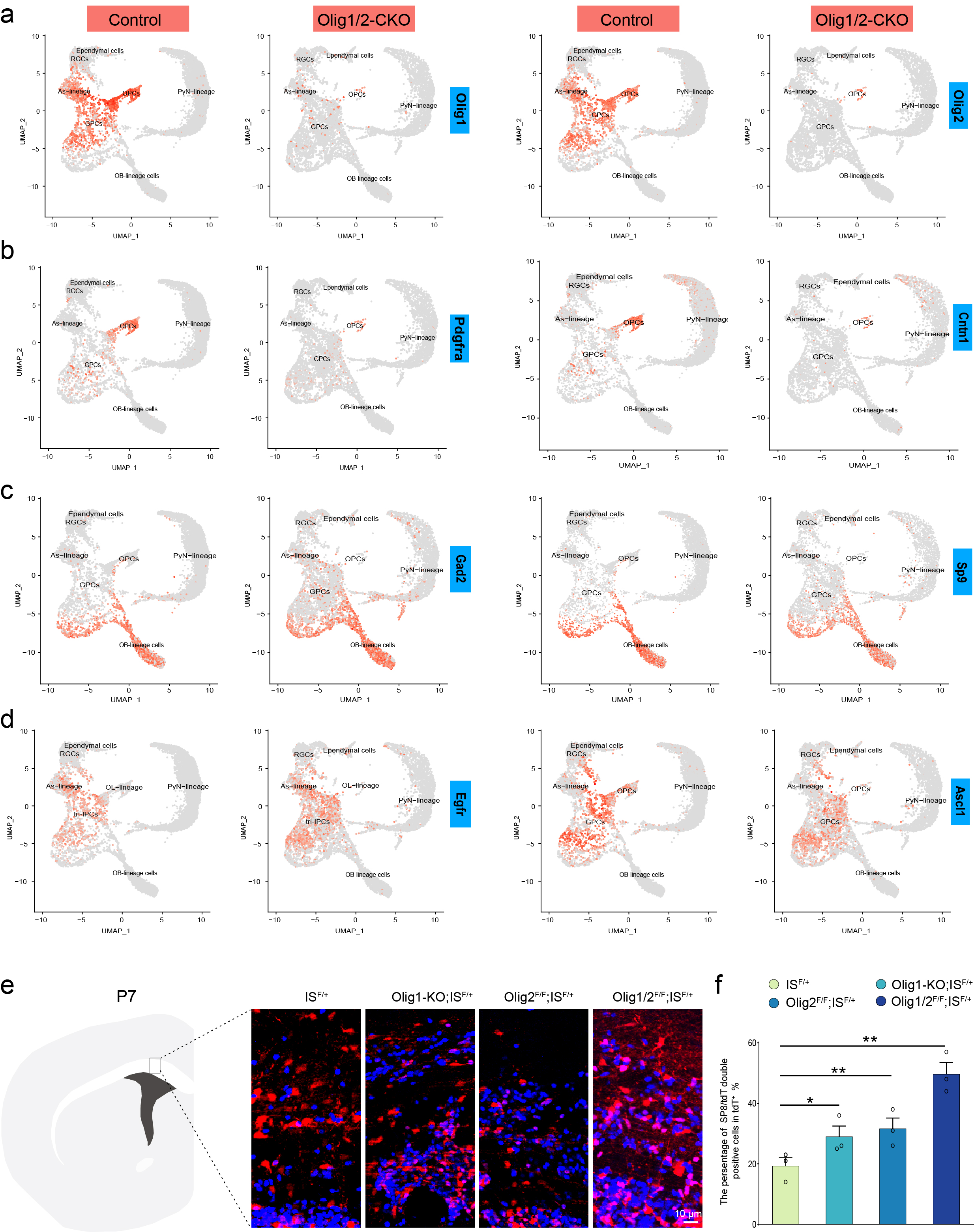

### Supplemental Figure4

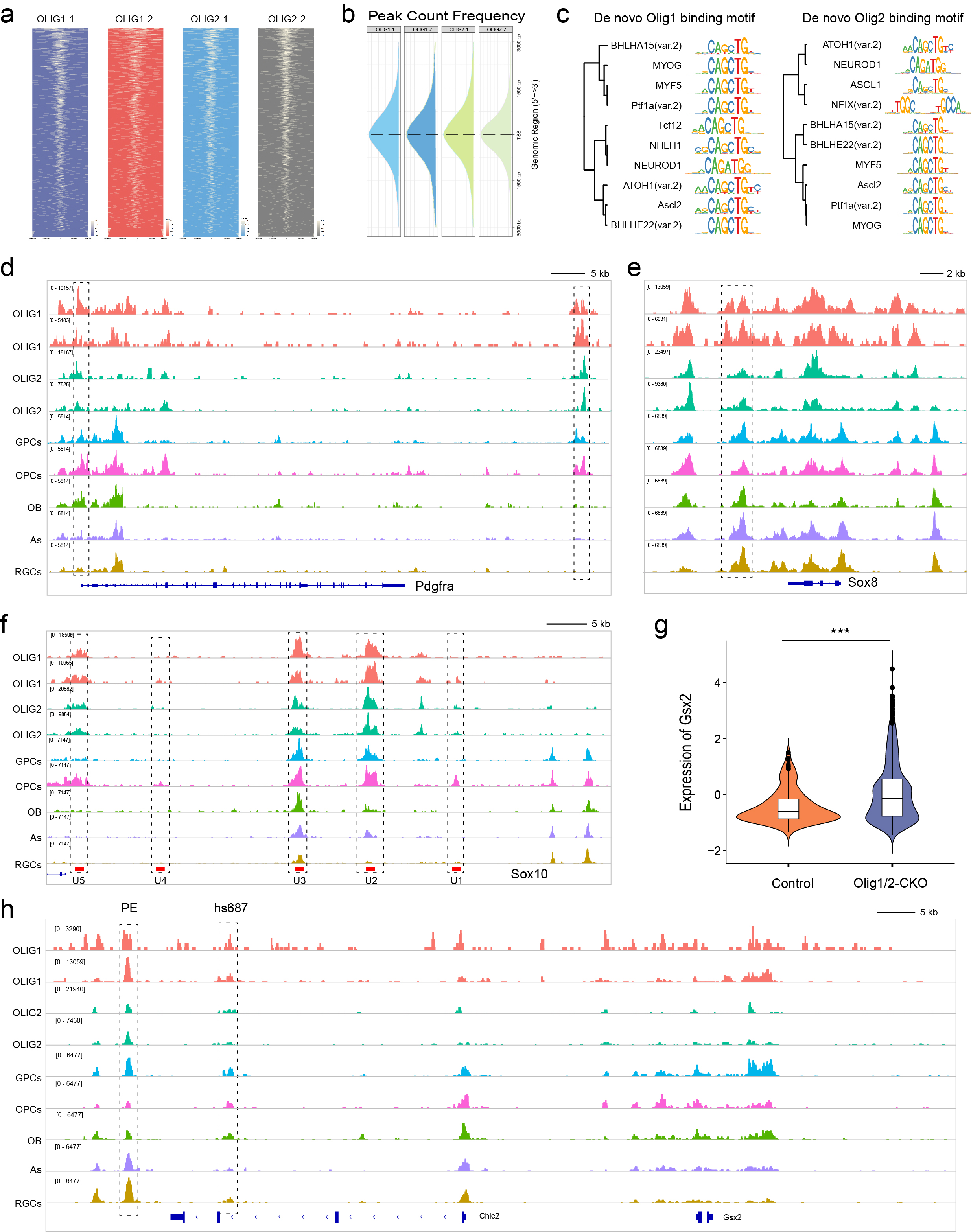

### Supplemental Figure5

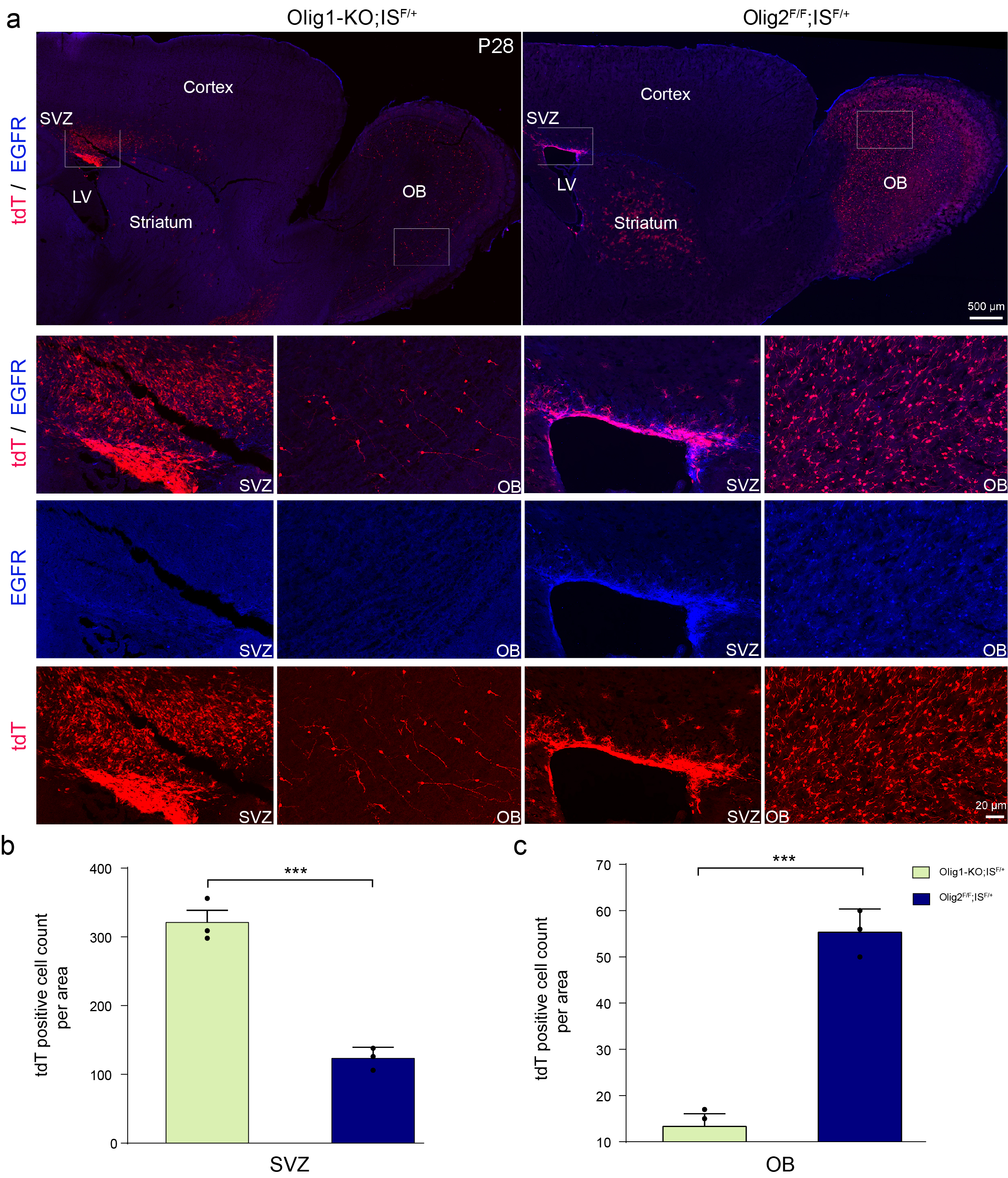

### Supplemental Figure6

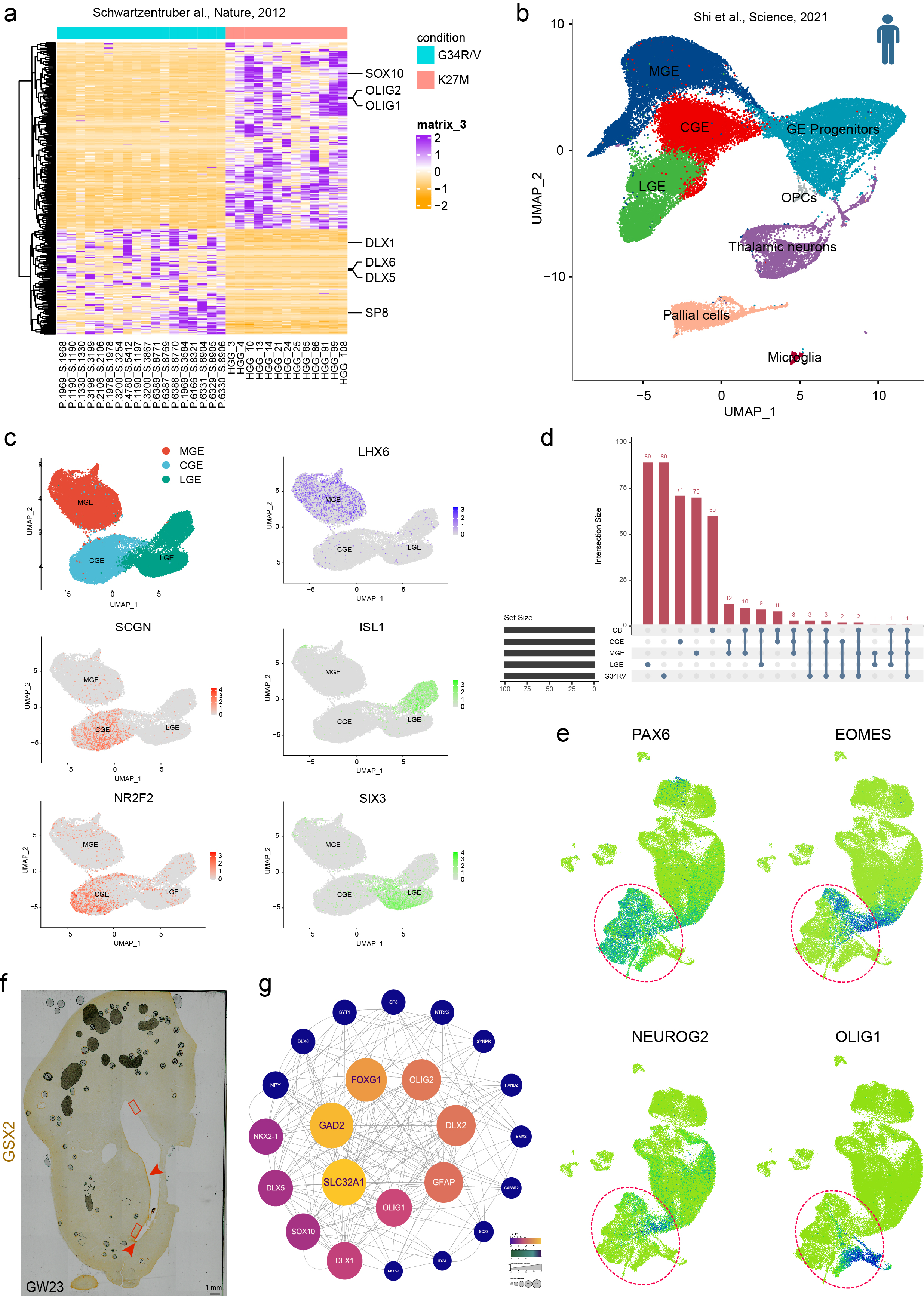
